# Prediction of Tubulin Inhibitor Sensitivity in Colon Cancer

**DOI:** 10.1101/2025.01.14.633026

**Authors:** Jose Liñares-Blanco, Lea Lemler, Irene Chicote, Javier Ros, Anna M Alcántara, Jordi Martínez-Quintanilla, Elena Elez, Héctor G Palmer, Carlos Fernandez-Lozano, Jose A Seoane

## Abstract

**Purpose:** Colorectal cancer is the second most common cause of cancer death worldwide. Despite advances in molecular strategies for stratifying patients with colorectal cancer, their clinical implementation remains limited. Previous efforts have mainly focused on the development of prognostic gene expression signatures; however predictive biomarkers remain essential for translating molecular insights into effective therapies and ultimately enhancing patient outcomes. This study advances the translation of a molecular signature identified in colon cancer toward new therapeutic options for clinical applications.

**Experimental Design:** We leveraged public gene expression datasets to develop a prognostic 37-gene signature in colon cancer and validated its prognostic value across six clinical cohorts. We subsequently investigated the biological processes associated with the signature and its predictive potential for therapeutic response. We further evaluated its predictive capability for tubulin inhibitor response using paclitaxel in vitro and in vivo in patient-derived preclinical models.

**Results:** The signature showed prognostic performance comparable to published gene expression signatures across six independent cohorts without clinical covariates. The signature was associated with the mitotic phase of the cell cycle and identified a population of patient-derived colon cancer models sensitive to tubulin inhibitor drugs, subsequently referred to as the Colon Tubulin-Inhibitor Expression Signature (ColoTubIn). Low ColoTubIn scores were associated with increased paclitaxel sensitivity.

**Conclusion:** Hence, ColoTubIn performs comparably to existing prognostic signatures while additionally identifying a molecularly defined subgroup of colon cancers with therapeutic potential. ColoTubIn is of particular interest since it can serve as a basis for drug repurposing strategies leveraging existing safety data to accelerate translation into the clinic.

**Translational Relevance:** Treatment options for colorectal cancer (CRC) remain limited, particularly following failure of first line chemotherapy. Given the high mortality associated with CRC, there is a pressing need for novel therapeutic strategies and predictive biomarkers to guide personalised treatment. We developed the Colon Tubulin Inhibitor Expression Signature (ColoTubIn), a transcriptomic signature that identifies a subset of patients with poor prognosis who may benefit from tubulin inhibitor therapy. Tubulin inhibitor Paclitaxel is routinely used in breast, lung, and ovarian cancers and has a well characterised safety, tolerability, and toxicity profile. Its repurposing in colon cancer could therefore facilitate clinical translation while reducing the time, cost, and risk associated with novel drug development. ColoTubIn provides a biomarker guided framework for patient selection in clinical trials evaluating paclitaxel and other tubulin inhibitors. Following prospective validation, this approach could expand therapeutic options and inform biomarker guided treatment strategies for patients with second line and refractory colon cancer.

## 1. Introduction

Colorectal cancer (CRC) is the third most commonly diagnosed cancer and the second leading cause of cancer-related deaths worldwide. Estimates for 2026 project over 150.000 new cases and approximately 55.000 deaths in the US^1^.

Traditionally CRC patients have been stratified based on histologic subtype, morphologic grade and American Joint Committee on Cancer (AJCC) Tumour, Node, Metastasis (TNM) stage to inform therapeutic approaches and prognostication^2,3^. In recent years new molecular biomarkers have been introduced, which reshaped treatment interventions strategies and prognostic outlook particularly in the metastatic setting ^4^: CRC tumours with high microsatellite instability (MSI) or mismatch repair deficiency (dMMR) have shown impressive responses to immune checkpoint inhibitors, which ultimately impact the overall survival of this subgroup. In addition, high MSI status has been associated with a favourable prognosis in stage II and can serve as adjuvant decision making tool ^5,6^. For patients harbouring a BRAF-V600E mutation, which is recognised as an unfavourable prognostic marker, targeted BRAF inhibition strategies have been developed for metastatic disease and demonstrated clinical benefit^7^. Moreover, RAS/BRAF wildtype tumours have been associated with better overall survival and RAS/BRAF mutation status has been shown to determine whether anti-EGFR therapy is a beneficial addition to standard chemotherapy in metastatic disease^8,9^.

However, besides the MSI status, no direct clinical use of the molecular subtypes in the non-metastatic setting has been found and they are therefore not routinely tested ^5^. Moreover, given the heterogeneous nature of CRC, studies in the metastatic setting have reported variable behaviour among the described biomarkers^10–12^. This highlights the urgent need for new prognostic and especially predictive biomarkers in CRC. Considering the high mortality rate of CRC, establishing new clinical standards is essential to ensure informed and personalised treatment decisions.

To date, several studies in colon and colorectal cancer have identified gene expression prognostic signatures ^13–25^ with two of them included in the National Comprehensive Cancer Network (NCCN) guidelines^26^. The primary goal of these signatures is to classify colon or colorectal cancer patients into risk groups, predict overall survival, disease recurrence or relapse. While some signatures provide predictive value for response to conventional chemotherapy, potentially helping to reduce unnecessary toxicities^20^, others lack real predictive validity due to the absence of a non-chemotherapy control groups as a baseline for comparison^15^. Hence, these signatures cannot truly establish whether a gene signature is predictive rather than simply prognostic identifying differences in tumour biology. Additional efforts have been made to forecast therapy response, however these signatures also failed to show true predictive capacity, are mainly limited to existing treatment options (e.g. chemotherapy, anti-EGFR target therapy, anti-angiogenic therapy)^27–31^ and novel therapeutic targets and treatment options remain mainly unexplored. Given the limited treatment options in CRC, particularly once first-line chemotherapy failed, new therapeutic targets are needed, which can offer new avenues for the management of CRC. In case new treatment options are identified^32^, existing gene expression signatures rarely progress beyond computational identification in the discovery dataset and confirmation in validation data set, lacking subsequent experimental validation^33^. In addition, it should be noted that discovery and validation datasets themselves often have inherent limitations due to their small sample sizes. Apart from this, many signatures are highly correlated, resulting in similar predictions across diverse signature sets. This can occur because they either neglect the adoption of novel feature selection methods that avoid the use of correlated gene features, thereby compromising predictive accuracy^34^, or they fail to sufficiently account for relevant prognostic clinicopathological variables^35^ or both. Moreover, although colon and rectal tumours have common characteristics, they should be defined separately due to differences in their molecular carcinogenesis, pathogenesis, and treatment^36^. While the described limitations capture only some of the weaknesses inherent in current gene signatures, they nonetheless illustrate potential reasons why these signatures have not achieved clinical implementation.

Hence, robust molecular signatures with both prognostic and predictive capacities are lacking. In this work, we bridge the gap between the discovery of a molecular signature in colon cancer, and its potential translation into novel therapeutic targets for patients in the clinic.

We present Colon Tubulin-Inhibitor Expression Signature (ColoTubIn), which is a signature developed via a multi-step pipeline to select prognostic genes in colon cancer patients from expression data.

ColoTubIn shows a higher accuracy than random gene sets, as well as 14 published colon and colorectal cancer-specific prognostic signatures in the internal validation. In the external validation, we compared ColoTubIn with the previously identified top six colon cancer signatures (mda114^15^, coloGuideEx^13^, coloGuidePro^14^, coloPrint^16^, Zhang^25^ and Immunologic Constant of Rejection (ICR)^37^) in six different external datasets. Apart from showing robust prognostic accuracy compared to current signatures, ColoTubIn exhibits significant predictive capability. High ColoTubIn signature score has been associated with resistance to tubulin inhibitor drugs, which was further validated with paclitaxel, a tubulin inhibitor, in vitro and in vivo in patient-derived preclinical models. Paclitaxel is routinely used in lung, breast and ovarian cancer treatment^38^. Given its established safety, tolerability, and side-effect profile, clinical translation could be achieved within a shorter timeframe and at reduced cost, while minimising clinical risk.

Hence, it presents a promising avenue for redefining future treatment paradigms in colon cancer for a subset of patients. Finally, ColoTubIn provides a framework for a straightforward evaluation of alternative therapeutic strategies incorporating additional tubulin inhibitors in the clinical setting.

## 2. Methods

### 2.1 Patients and tumour samples

ColoTubIn was developed using a retrospective cohort of 325 patients with primary colon adenocarcinoma (COAD) collected between 2009 and 2015 from multiple tissue source sites worldwide through The Cancer Genome Atlas (TCGA) program ^39^. As described in the foundational TCGA colorectal study, eligible samples were obtained from primary surgical resections of patients who had received no prior chemotherapy or radiotherapy for their disease^40^. As TCGA was a multi-center tumour characterisation project rather than a clinical trial, treatment was not standardised across participating institutions and followed local standards of care ^39^. Fresh frozen primary tumour tissue was collected across surgical stages and histologic grades ^40^. RNA was extracted from tumour specimens using a modified RNA AllPrep protocol (Qiagen), as previously described^40^.

Six independent cohorts comprising primary colon tumour samples were included for validation. The E-MTAB-12862 cohort comprised samples collected at Uppsala University Hospital and Umeå University Hospital ^41^. Samples from the UK-wide Stratification in Colorectal Cancer (S:CORT) consortium were obtained from the SPINAL sub-cohort^42^ and ^43^. The remaining validation cohorts were GSE17536 ^44^, GSE17537 ^44^, GSE29621 ^45^ and GSE39582 ^46^.

Comprehensive descriptions of the individual validation cohorts can be found in the corresponding publications, with patient characteristics of the discovery and validation cohorts summarised in Supplementary Table S2.

This study adheres to the REMARK guidelines for reporting prognostic biomarker studies ^47^, details can be found in Supplementary Table S3.

### 2.2 Data acquisition and preprocessing

The RNA-Sequencing (RNA-Seq) data (Fragments Per Kilobase of transcript per Million mapped reads – Upper Quartile normalized (FPKM-UP)) of colon primary solid tumour samples were downloaded from The Cancer Genome Atlas (TCGA) repository using the TCGABiolinks (version 2.17.1) R package^48^. Protein coding genes were selected using the biomaRt (version 2.44.0) R package. Genes with an FPKM count of zero in more than 20% of the samples were removed. In order to perform future external validation, the genes available in TCGA were intersected with the genes available in the Affymetrix Human Genome U133 Plus 2.0 microarray platform (GPL570). Finally, a total of 17,267 genes were defined as the search space of the gene selection algorithm.

Clinical annotated data, such as stage, sex, MSI status, molecular subtype, and age at diagnosis, were obtained^49^. Only samples with available annotated overall survival times and events were included. Overall survival was defined as the time from diagnosis to death from any cause. Categorical clinical variables were transformed into dummy variables. Duplicated samples from the same patients were selected based on the highest mean expression profile, resulting in 325 patients from the TCGA Colon adenocarcinoma (TCGA-COAD) cohort.

Variance Stabilizing Transformation (VST) with DESeq2 (version 1.28.1) R package was calculated for each gene ^50^. Ten percent of the samples were excluded as the test set (33 samples), whereas the remaining 90% of the samples were used as train set (292).

In total, six validation cohorts containing primary colon tumour samples from the RNA-Seq cohort EMTAB-12862 as well as five microarray cohorts were defined.

The RNA-Seq data of cohort EMTAB-12862 of colon primary tumour samples was obtained from ArrayExpress under accession number E-MTAB-12862^51^. Gene expression was quantified as transcripts per million (TPM) and log2-transformed. A pseudo count of one was added to the TPM to avoid negative values for expression levels.

The microarray cohorts GSE17536, GSE17537, GSE29621 and GSE39582 were generated using Affymetrix Human Genome U133 Plus 2.0 Array and were downloaded from Gene Expression Omnibus (GEO). The remaining validation cohort SPINAL was based on Affymetrix Almac Xcel Array and was retrieved from SCORT database upon successful data request approval (https://www.s-cort.org/index.html). For all microarray-based validation cohorts the custom Chip Definition File (CDF) file for each platform was used to map the Affymetrix data to Ensembl gene annotations^52^. In addition, the Robust Multi-array Average (RMA) was applied for normalisation using the affy^53^ package (Version 1.82.0).

Clinical variables were retrieved when available. Sex and stage were collected for all external cohorts, while age was unavailable for GSE29621. As MSI status was inconsistently reported, missing MSI status was predicted using the R package PreMSIm (v1.0), which infers MSI from a 15 gene expression panel^54^. Samples lacking overall survival time or event information, as well as untreated samples from GSE39582, were excluded. For consistency, follow up was restricted to a maximum of 1,825 days, corresponding to five years, in all validation cohorts.

Given the heterogeneity of the datasets, the ComBat function, implemented in the R sva (version 3.36.0) package^55^, was used. In our case, ComBat function with parametric empirical Bayes frameworks for adjusting data for batch effects was applied. The TCGA-COAD train data was defined as the reference batch. Supplementary Figure S1 shows the adjustment of the batch effect according to the validation and training cohorts.

### 2.3 Feature selection method for ColoTubIn gene selection

In this work, a predominant correlation analysis^56^ was conducted to evaluate gene correlation in colon adenocarcinoma expression data and to filter the most informative genes, reducing the dimensionality of the analysis (Supplementary Figure S2). This approach is a multivariate filtering method, which uses the measure of entropy (H) and the Information Gain (IG) to search for the subgroup of dominant genes for a specific medical condition. The action of these two measures is encapsulated in the Symmetrical Uncertainty (SU). Defined as follows: First, the SU value was calculated for each gene, keeping relevant genes based on a threshold (0.025) and sorting them in descending order according to this value. Second, genes that provide redundant information were removed. The event of death in overall survival follow-up was used as a dependent variable. Consequently, we adopted a model-independent approach to select genes that exhibited a strong correlation with cancer survival, while minimising their correlation with uninformative genes (predominant correlation). In our investigation, this procedure was applied to the complete transcriptomic profile of each patient, encompassing a total of 17,267 genes. Among these, the algorithm identified 37 genes that met the specified criteria.

### 2.4 Internal validation of the ColoTubIn 37-gene signature

In order to determine whether the signature discriminates between two survival groups without bias, an internal validation was run with the R package SigCheck^57^ (version 2.20.0). The performance of SigCheck was calculated through a Support Vector Machine (SVM) classifier trained on the 90% training set and tested on the 10% test set, which had the same sample distribution as in the signature discovery, in four different scenarios. In each check a background performance distribution is calculated, and the performance of the signature is compared by calculating an empirical p-value. First, the sigCheckRandom function includes p-values calculated using random signatures of the same length (in this case, 37 genes). Second, the sigCheckKnown function compares the performance of the signature with a set of previously identified signatures. In our case, we used the whole The Molecular Signatures Database (MSigDB) database^58^, and also identified 14 signatures for colon cancer prognosis. Third, the test compares the performance of the signature on the dataset by performing permutations on the same dataset (N=10,000), namely, by permuting the survival class metadata. Finally, the fourth test randomly permutes the gene data, reassigning expression values for each gene across the samples. Both permutations are performed independently by running the sigCheckPermuted function. The top six signatures from the literature were used as a reference in the following analyses to compare our signature.

### 2.5 Construction and validation of the signature-based Cox Proportional Hazard model

In order to test the predictive performance of our 37-gene signatures, we constructed a Cox Proportional Hazard (CPH) model with and without clinical variables, including age, sex, stage, Chromosomal Instability (CIN) and MSI in the train set of TCGA-COAD. This resulted in three models, including i) the genes and the clinical variables, ii) only the genes, and iii) only the clinical variables constructed in the train set from TCGA-COAD. The three models trained in the TCGA-COAD train set were subsequently tested in the TCGA-COAD test set and the six external cohorts. The performance improvement of our model was compared against the top six signatures identified by the SigCheck package. A CPH model was created by each signature in the TCGA - COAD train set, following the same methodology. To account for the distinct censoring distributions within the external cohorts, we utilised Uno-consistent concordance index (C-index) for model evaluation (R package survC1 version 1.0-3), as it provides a more robust estimate of concordance than the traditional Harrell’s C-index used in the TCGA-COAD test, when censoring patterns vary^59^.

### 2.6 Patient-specific signature activity

We aimed to infer the activity of specific gene signatures using gene expression data from the TCGA-COAD cohort. To achieve this goal, we employed the GSVA R package^60^, specifically utilising the Pathway level analysis of gene expression (PLAGE) scoring method to calculate signature activities for each patient. We computed the activity of the ColoTubIn signature and applied the Mclust algorithm^61^ to assess whether distinct, independent distributions existed within the cohort, based on these activity values. The results of Mclust revealed the presence of two clusters, categorized as high and low PLAGE score. These cluster assignments were then utilised to relabel the patients within the cohort, facilitating downstream differential expression (DE) analysis between the two groups. Subsequently, we performed pathway enrichment analysis using the HTSanalyzeR2 R package^62^ to gain insights into the biological processes associated with these differential genes.

Given the enrichment of cell cycle pathways in our initial analysis, we pursued a more detailed investigation. We downloaded gene signatures corresponding to various phases of the cell cycle from the Reactome database. For each cell cycle phase, we calculated its activity and explored the relationships between these phases by computing Pearson correlations between pairs of signature-cell cycle phase combinations. This comprehensive methodology allowed us to investigate signature activity, identify distinct patient clusters based on plage scores, perform differential expression and pathway enrichment analyses, and delve deeper into the dynamics of cell cycle pathways within the TCGA-COAD cohort.

### 2.7 Drug-set enrichment analysis

We obtained gene expression data for colorectal cancer cell lines from the CCLE 21q2 dataset, which comprises 27 unique cell lines.

To mitigate potential batch effects between the Cancer Cell Line Encyclopedia (CCLE) dataset and TCGA cohort we applied the Combat algorithm from sva R package^55^. This step ensured that any variations due to dataset origin were removed before the subsequent analysis of gene signatures.

To evaluate the drug repurposing potential of specific genes, we employed the ColoTubIn probability-score. The score was calculated by fitting a CPH to the expression of signature genes in the train set of TCGA and obtaining linear predictions from the CCLE dataset.

We also acquired drug-cell line response data from the PRISM repurposing dataset (v19q4). This dataset included the Area Under the Curve (AUC) values for each drug-cell line combination after treatment, enabling us to assess the efficacy of drugs in CRC cell lines.

We calculated the correlation between the ColoTubIn probability-score and the AUC for each drug. Since we hypothesized that drugs with similar mechanisms of action (MoA) would have comparable effects on similar cell lines, we conducted a gene set enrichment analysis (GSEA) - like enrichment analysis based on the MoA, using the fgsea R package^63^. Instead of pathways we used the MoA of the drugs, and instead of genes we set the name of the drugs. This approach aimed to identify associations between drugs, drug types, and their effects on CRC cell line viability.

### 2.8 Patient-derived xenografts establishment

Patient-derived xenografts (PDX) from Héctor G. Palmer’s laboratory^64^ were used for preclinical validation of the gene expression signature. Briefly, human colorectal carcinoma tissues were obtained upon surgery in accordance with the ethical standards of the institutional committee on human experimentation. Histological diagnosis was based on the microscopic features of carcinoma cells determining the histological type and grade. The tumour tissues were washed 3 times in cold PBS solution and incubated overnight in DMEM/F12 (Gibco; Thermo Fisher Scientific) containing a cocktail of antibiotics and antifungals (penicillin (250 U/ml), streptomycin (250 mg/mL), fungizone (10 mg/mL), kanamycin (10 mg/mL), gentamycin (50 mg/mL), and nystatin (5 mg/mL; Sigma-Aldrich). Enzymatic digestion was performed using collagenase (1.5 mg/mL; Sigma-Aldrich; #C0130) and DNase I (20 mg/mL; Sigma-Aldrich; #D4263) in a medium supplemented with a cocktail of antibiotics and antifungals (penicillin (250 units/ml), streptomycin (250 mg/mL), fungizone (10 mg/mL), kanamycin (10 mg/mL), gentamycin (50 mg/mL), and nystatin (5 mg/mL)) for 1 hour at 37°C with intermittent pipetting every 15 minutes to disperse the cells. The dissociated sample was then filtered (100 mm pore size) and washed with fresh medium. Red blood cells were lysed by 10 minutes exposure to ammonium chloride and the sample was washed again. Finally, 1x105 cells were resuspended in 50 µLof PBS and mixed with 50 µLof Matrigel, and subcutaneously injected into NOD-SCID flank mice to generate subcutaneous tumours for further histological and RNA analysis.

### 2.9 Colorectal Patient-derived xenografts sequencing

RNA was isolated from 80 fresh frozen colon PDXs, using Rneasy Mini Kit (QIAGEN). Each sample was quantified using NanoDrop. RNA-Seq libraries were prepared and paired-end 150 bp sequencing (50M reads per sample) was performed on a NovaSeq 6000s (Illumina) in accordance with the manufacturer’s instructions. Counts were obtained using the Next-flow RNA-seq pipeline version 3.8.1 (https://nf-co.re/rnaseq/3.8.1), using GRCh38 genome and gencode V41, STAR for annotation, and RSEM for quantification.

### 2.10 Patient-derived xenografts organoid model generation

The PDXs with the five highest and five lowest ColoTubIn probability-score were selected for PDX organoid (PDXO) model generation, including T138 (*KRAS*G12V and *BRAF*WT), CTAX23 (*KRAS*WT and *BRAF*V600E), CTAX26 (*KRAS*G12D and *BRAF*WT) and T224 (*<u>KRAS</u>*WT and *BRAF*V600E), T245 (*KRAS*WT and *BRAF*V600E), T251 (*KRAS*WT and *BRAF*WT). The selected subcutaneous CRC-PDX tumours were dissociated with collagenase and DNase I for 30 minutes at 37°C. The cellular pellet was mixed with Matrigel and plated in 6 well plates with human intestinal stem cell media (HISC adapted from^65^) complemented with Rhock and a GSK3 inhibitor to generate PDXO cultures. The HISC media was prepared with 28% Wnt3A conditioned medium + 14% RSPO1 conditioned medium + 7% noggin conditioned medium + basal culture medium supplemented with B27 (1X; Gibco, 17504044), N2 (1X; Gibco, 17502048), N-Acetyl-L-Cysteine (1,35 mM; Sigma-Aldrich, A9165-5G), Nicotinamide (5mM; Sigma-Aldrich, N3376), human EGF (50ng/ml; PrepoTech, AF-100-15), [Leu15]-Gastrin I (10 nM; Sigma-Aldrich, G9145-1mg), Prostaglandin E2 (10 nM; Sigma-Aldrich, P0409), A83-01 (500 nM; Tocris, 2939), SB202190 (2.8 µM; Sigma-Aldrich, S7067) and Normocin (125µg/ml; Invivogen, ant-nr-2). Conditioned medias were obtained from confluent L1wnt3A cell line (Wnt3A conditioned medium), 293t RSPO-mCherry cell line (RSPO1 conditioned medium), and 293t Noggin-mCherry cell line (noggin conditioned medium) cultured in basal media.

### 2.11 Colon PDXO model treatment with paclitaxel in vitro

Confluent organoid cultures were incubated with ice-cold cell recovery solution (Corning, 354253) at 4°C for 30 minutes following the manufacturer’s protocol. Organoids were collected, washed with basal media, mixed with 200-400 µl TrypLE (Gibco, 12605010), and incubated at 37°C for 2-3 minutes. The cell suspension was mixed up and down by pipetting, washed with basal media, and counted using a Neubauer chamber.

Single cell organoid suspension (2.5x103-5x103) from T138, T224, T245, T251, CTAX23, and CTAX26 PDXO models were plated in a 5 µl drop of Matrigel (cells diluted in basal media and mixed with Matrigel in a proportion 1:4) in a 96-well plate and cultured with HISC media for 48 hours. Then, cells were prepared via 1:2 serial dilutions of paclitaxel (19 doses starting from 100 µM) or with a vehicle control in DMEM 2.5% FBS. Treatment refreshment was performed at 72 hours. CellTiter-Glo (Promega, G7571) was used (following the manufacturer’s protocol) at 5 days to determine cell viability.

### 2.12 Study Mice preparation

Experiments with mice were conducted following the European Union’s animal care directive (86/609/CEE) and were approved by the Ethical Committee of Animal Experimentation of the VHIR - the Vall d’Hebron Research Institute (approval ID, 17/15 CEEA, 18/15 CEEA and 12/18 CEEA). All the animals used in this study were female, (NOD.CB17-Prkdcscid/scid/Rj) aged between 6-8 weeks (21-24g), and were purchased from Janvier Laboratories. Subcutaneous PDX tumours were generated by mixing 1x105 tumour cells resuspended in 50 µl of PBS with 50 µl of Matrigel and injecting them subcutaneously into a flank of a NOD-SCID. Tumours were measured every 2-3 days with a caliper until they reached 100-200mm3, when mice were randomized by tumour size, and those that died before the end of the experiments were excluded. The experiments were not performed in a blinded fashion. All the mice were closely monitored by the authors, facility technicians, and a veterinary scientist responsible for animal welfare. The mice were maintained in a specific-pathogen-free facility under controlled temperature and humidity and given ad libitum access to a standard diet and water.

### 2.13 Subcutaneous tumour xenografts

The experimental groups were treated with either vehicle (PBS) or Paclitaxel (7,5 mg/kg twice per week) by intraperitoneal injection. When matching endpoint criteria of 1cm3 of tumour volume, mice were euthanized, and parts of the xenograft tumours were snap-frozen or fixed in paraformaldehyde for histological analysis. Tumour growth was monitored by caliper measurement three times per week, and tumour volume was estimated using the following formula: V = (length x width2)/2, where length represents the largest tumour diameter and width represents.

## 3. Results

In this study a 37-gene prognostic as well as predictive signature for colon cancer, ColoTubIn, was discovered. Genes within ColoTubIn are enriched for pathways related to the mitotic phase of the cell cycle. The signature revealed a subgroup of colon cancer patients with sensitivity to tubulin-targeting agents. The therapeutic relevance of this finding was demonstrated in vitro and vivo with a commonly used tubulin inhibitor, which showed efficacy in patient-derived preclinical models for patients with low ColoTubIn signature score.

### 3.1 Development of a transcriptional signature associated with prognosis in colon cancer

A feature selection algorithm was used to select genes with high correlation with patient outcome, but low correlation among them. By employing 90% of TCGA-COAD (N=292) as the training data, we selected 37 genes, the ColoTubIn signature (**Figure 1a and b**). The 37 genes, combined with clinical variables, were used to generate a multivariate Cox Proportional Hazard model for overall survival in the training dataset (**Figure 2a).** Four clinical variables (Age at diagnosis,

**Figure 1:**
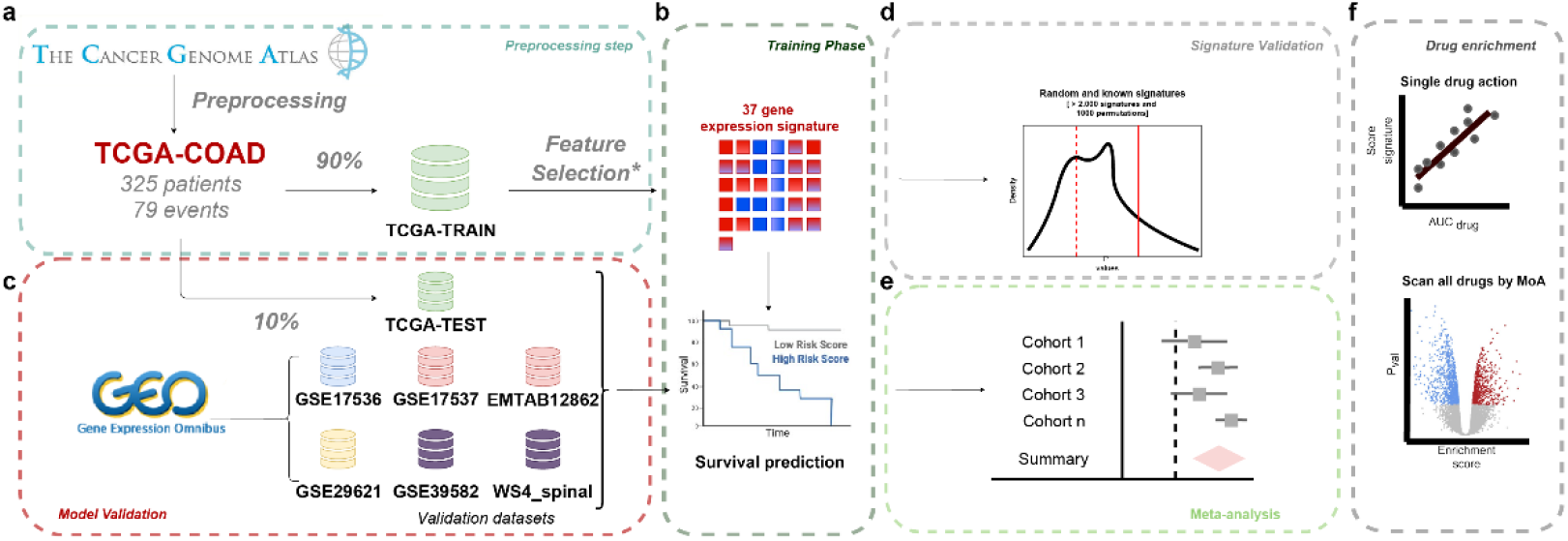
Scheme of the methodology used to generate and validate the ColoTubIn signature: a) We applied feature selection over 90% of the TCGA-COAD dataset. b) A Cox Proportional Hazard model was trained in the training dataset and c) validated in 10% of the TCGA cohort and the other seven independent cohorts. d) The SigCheck package was used to compare the selected features with random combinations of features. e) We compared the results of our signature with those of other relevant CRC signatures. f) We evaluated whether our signature score is predictive in cell line drug resistance datasets.

**Figure 2:**
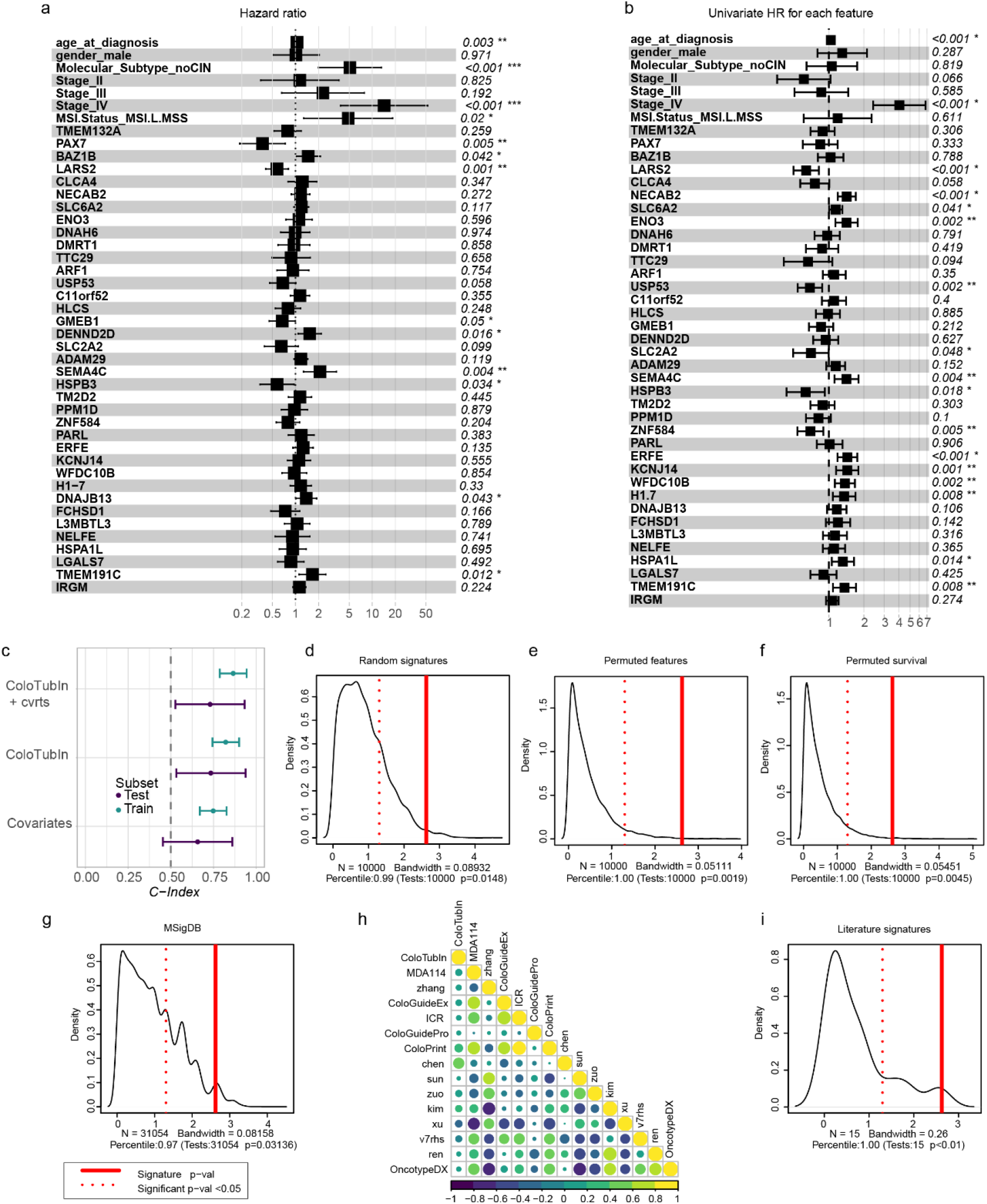
Evaluation of the Cox Proportional Hazard Model. Panels a) shows the multivariate hazard risk of all the features included in a Cox Proportional Hazard model. The forest plot shows the hazard ratios and p-values for each gene and covariate; Panel b) represents the univariate hazard risk of all the features included in a Cox Proportional Hazard model. Panel c) shows the C-index in train (n=292) and test (n=33) data. To infer gene effect in the model, C-index were calculated in three different conditions in the training phase (with genes, clinical covariates, and both combined). Panel d) Density plot representing SigCheck (association with overall survival) evaluations of ColoTubIn compared with 10,000 random gene signatures of the same size. Panel e) shows the comparison of ColoTubIn vs 10,000 permutations of the genes. Panel f) represents ColoTubIn vs 10,000 permutations over survival events. Panel g) shows the comparison between ColoTubIn and all signatures in the MSigDB database. Panel h) represents the correlation between gene set variation analysis scores of ColoTubIn and 14 literature signatures associated with colon cancer outcome. Panel i) represents the comparison between the 14 literature signatures associated with colon & colorectal cancer outcome and ColoTubIn. In each SigCheck plot, continuous red lines represent the p-value achieved by the ColoTubIn signature, whereas dotted red lines represent values at which the p-value is significant.

Molecular Subtype, Stage IV, and MSI Status) and nine genes (*HSPB3, SEMA4C, LARS2, GMEB1, DENND2D, DNAJB13, PAX7, BAZ1B, and TMEM191C*) were significantly associated with overall survival in the multivariate model. Similar trends were identified in the univariate analyses (**Figure 2b**).

The additive prediction of the ColoTubIn signature was tested by generating three CPH models, including i) only clinical variables, ii) only the 37 genes, or iii) both. In training, the model including genes and clinical variables reached the highest C-index and AUC (C-Index=0.86; AUC=0.88), whereas the model trained with only clinical variables scored the worst (C-Index=0.74; AUC=0.77), and the model trained with only genes performed close to the combined model (C-Index=0.82; AUC=0.85). In the test set (n=33, 10% TCGA-COAD), the model with the worst performance was the clinical variables model (C-Index=0.65; AUC=0.60), whereas the model with the best performance was the gene-only model (C-Index=0.73; AUC=0.65), and the model with genes and clinical variables achieved similar results to the gene-only model (C-Index=0.72; AUC=0.63) (**Figure 2c**). These results in the TCGA cohort indicate that the addition of the ColoTubIn gene list to a clinical variable model provides superior prognostic performance compared with the use of only clinical variables.

In order to evaluate whether the ColoTubIn signature achieves better overall survival prediction than randomly generated signatures, we performed multiple analyses in the TCGA COAD cohort (Figure 1d). First, we compared whether the ColoTubIn signature performed better than 10,000 random signatures with 37 genes using SigCheck^39^ (**Figure 2d**). The ColoTubIn signature was within the top 1% of the gene sets (p=0.0148), indicating that the gene list was not randomly associated with the outcome. The second and third scenarios compared the ColoTubIn signature with 10,000 permutations of the samples across the genes (**Figure 2e**) or across the outcome (**Figure 2f**). The comparison of ColoTubIn with a random permutation of samples across the genes (p=0.0019) and across the outcome (p=0.0045), which ranked first in both cases, revealed that there was no sampling bias in the development of the signature. Finally, in the fourth scenario, we compared ColoTubIn with all MSigDB signatures (N=31,054), ranking among the top 3% of all signatures (p=0.031, **Figure 2g**). Moreover, the prognostic performance of ColoTubIn and the 14 signatures was compared within SigCheck. The ColoTubIn signature achieved the highest performance in predicting overall survival (p<0.01, **Figure 2i**), followed by the next top six performers: MDA114, zhang, coloGuideEx, coloPrint, coloGuidePro and ICR. These top six signatures were also evaluated using the first three scenarios via the SigCheck package (random gene selection and both permutation comparisons). Unlike ColoTubIn, these signatures do not outperform random models; only MDA114 and zhang reached statistical significance when tested against permuted features and outcome in TCGA **(Figure S3)**.

Finally, possible correlations between ColoTubIn and 14 colon and colorectal cancer prognostic signatures were investigated. ColoTubIn score exhibits a very low correlation with other colon and colorectal prognostic signatures **(Figure 2i)**, whereas most of the other signature scores were highly correlated with each other. These results revealed that ColoTubIn uses different genomic information than the other signatures.

### 3.2 Validation of the ColoTubIn signature

Next, we validated ColoTubIn in external cohorts (**Figure 1e)**. We compared ColoTubIn signature against six colon cancer signatures with the top performance (MDA114, ColoGuideEx, ColoGuidePro, ColoPrint, Zhang and ICR) following the SigCheck experiment (see Figure 2i). In addition, a signature based solely on clinical variables was incorporated to quantify the added predictive value of the transcriptomic signatures in the comparative analysis. Samples without overall survival time, event, or non-treated samples (GSE39582) were removed. For each gene signature, we generated Cox Proportional Hazard models with and without covariates. The Uno C-index for each model per validation cohort is shown in the figure **(Figure 3a)**. ColoTubIn predictions using genes and covariates shows a baseline performance compared with average, and performing best in the in the GSE39582 cohort (C-index = 0.72 [0.65-0.81]). In the meta-analyses ColoTubIn achieved a C-index of 0.67 ([0.63-0.72]). Notably, none of the models performed consistently better than covariates alone. When restricted to gene expression predictors, ColoTubIn achieved the best accuracy compared to other signatures in the cohorts GSE29621, GSE39582, TCGA test and SPINAL. Overall, ColoTubIn obtained a C-index = 0.58 [0.54-0.62]).

**Figure 3:**
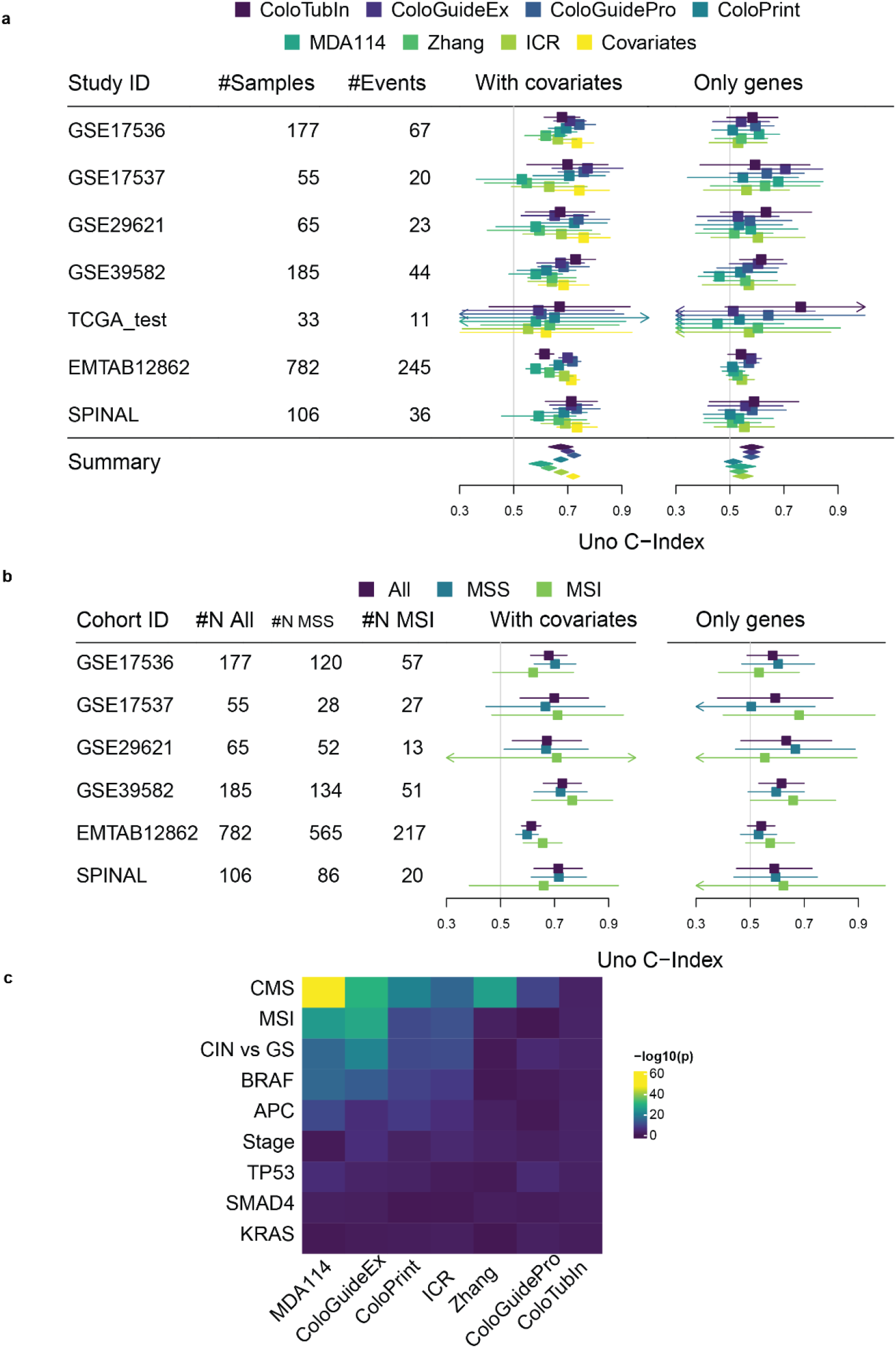
External validation of ColoTubIn and comparison with the top colon cancer signatures. In Panel a) we show the C-index achieved for each signature. These Cox Proportional Hazard models were built with gene signatures information and clinical information or with gene expression alone. In addition, a Cox Proportional Hazard model (in yellow) with only clinical information was built in order to compare performance. In Panel b), we show the C-index from Cox Proportional Hazard models comparing MSI vs MSS samples with gene expression and covariates and with gene expression alone. Panel c) shows p-values from statistical tests performed by comparing enrichment scores for each signature with clinical variables. The Wilcoxon test was performed on variables with two groups, whereas the Kruskal-Wallis test was used for those with three or more groups.

Next, to ensure that ColoTubIn is not related to MSI in colon cancer, we generated the ColoTubIn CPH model independently by each MSI/microsatellite stable (MSS) patient subgroup. The results did not reveal significant differences between MSI status using gene expression and clinical variables or gene expression alone (**Figure 3b**).

### 3.3 The Cell Cycle Mitotic Phase drives ColoTubIn signature

In order to identify the biological processes behind the ColoTubIn signature, we calculated a single-sample enrichment score for ColoTubIn and the top six signatures. We compared these scores with clinical variables and molecular biomarkers of colon cancer (including MSI, consensus molecular subtypes (CMS)^66^, CIN vs Genomically Stable (GS) subtype, or *BRAF*, TP53, APC, SMAD4) to test the independence of the signatures from these factors **(Figure 3c and Supplementary Table S1)**. The results indicate that the CMS is the variable most strongly associated with the signatures (5.57e-10<=p<=1.80e-48), whereas ColoTubIn has a relatively low association (p=6.14e-3). Among the other variables, large effect sizes were observed for the molecular subtype, MSI, *BRAF*, and *APC*. However, ColoTubIn showed a weaker correlation with the colon biomarkers in comparison to the other signatures suggesting that its performance is less affected by them.

The distribution of enrichment scores of the ColoTubIn signature does not follow a normal distribution, with an unexpected number of patients having elevated values (**Supplementary Figure S4a**). We conducted a model-based clustering based on finite normal mixture modeling to analyse whether this may be caused by different molecular characteristics across patients with high or low ColoTubIn score. Supporting our hypothesis, the analysis identified two different clusters, spotting a subgroup in high ColoTubIn score patients.

A differential expression analysis was subsequently performed between these two patient groups (high vs. low ColoTubIn score) followed by a gene set enrichment analysis employing the Reactome database (**Figure 4a**). We identified several pathways related to the cell cycle, including the most enriched *pathway establishment of sister chromatid cohesion* (**Figure 4b**) and the *mitotic prometaphase* pathway (**Figure 4c**). Enrichment plots are shown also for the *M-phase* **(Figure S4b)** and *Cell cycle mitotic pathways* (**Figure S4c**). These results indicate high cell cycle dysregulation in patients with higher ColoTubIn score, specifically in the mitotic phase.

**Figure 4:**
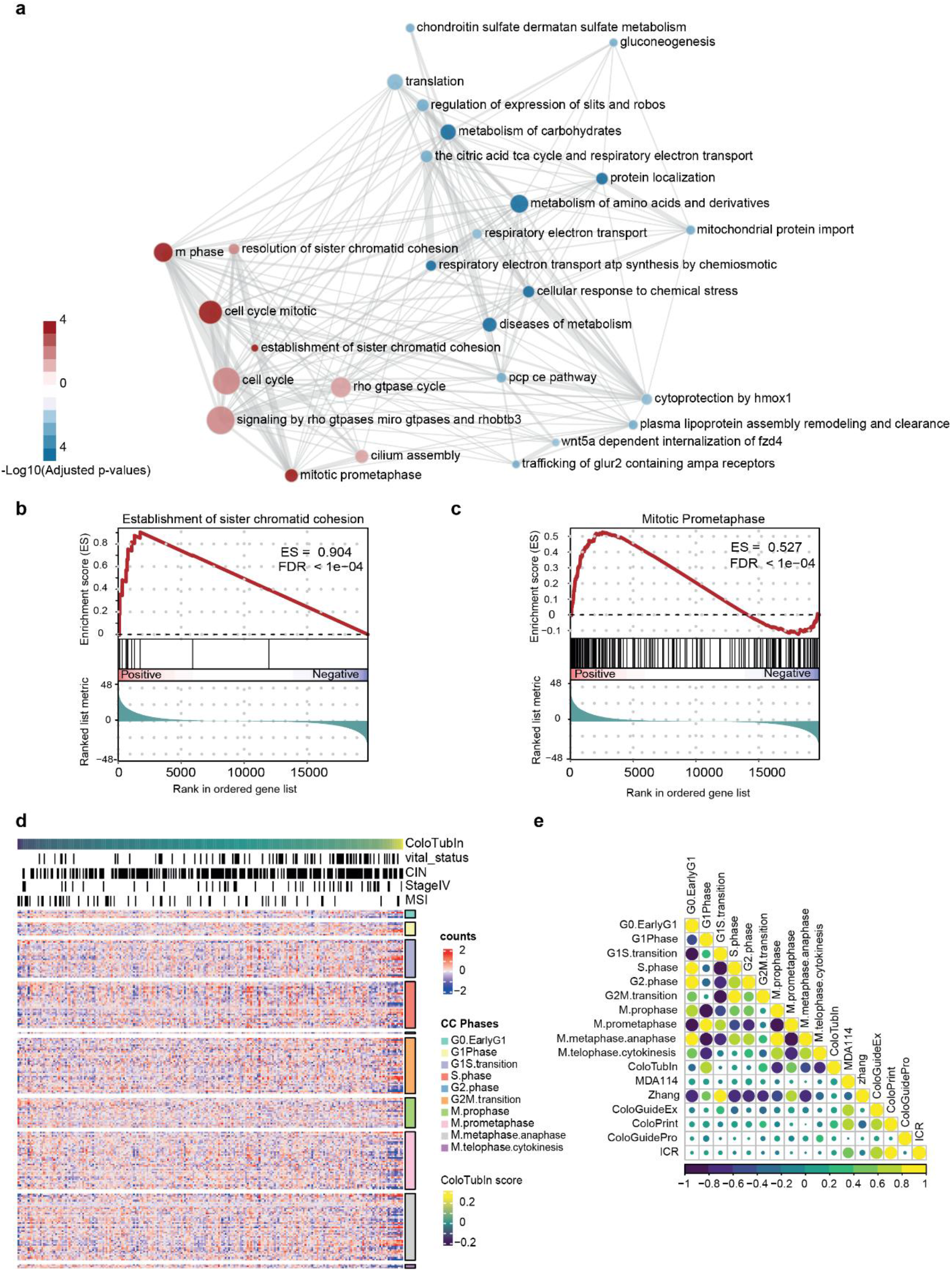
ColoTubIn is associated with Cell Cycle Mitotic Phase. Panel a) shows the results of the pathway enrichment of ColoTubIn high (N=17) vs ColoTubIn low (N=308) samples. The colours indicate significance (-log10 adjusted p-value) and direction (positive enrichment red, negative enrichment blue) and the size of the circle represents the size of the pathway. Shared genes across the pathways are represented by the thickness of the edge. Panels b) and c) represent the GSEA plots of the two main hits of the pathway enrichment analysis, “Establishment of sister chromatid cohesion” and “Mitotic Prometaphase”, respectively. The heatmap in panel d) represents in the rows the gene expression of all the genes related to the cell cycle, grouped by cell cycle phases, and in the columns, the samples from TCGA-COAD were sorted by the ColoTubIn signature. Panel e) measures the correlation across the signatures PLAGE-scores and the cell cycle phases, with yellow being positively highly correlated and blue being negatively highly correlated. The size of the circle represents the significance of the correlation.

To elucidate the precise relationship between ColoTubIn and the cell cycle, we analysed the expression patterns of genes implicated in each phase of the cell cycle. From the Reactome database, we extracted the genes involved in ten different phases of the cell cycle. The expression of these genes per phase in TCGA-COAD patients is shown using a heatmap in which patients are sorted by the ColoTubIn score (**Figure 4d**). We observed highly similar gene expression patterns in the cell cycle gene sets of patients with high ColoTubIn score.

Next, we calculated the pathway activity scores per patient for each cell cycle phase gene-set and tested the correlation between and with the ColoTubIn scores (**Figure 4e**). We observed that the ColoTubIn score is highly correlated with the scores of all the M-phases of the cell cycle and, to a lesser extent, with the G1-phase score. The major families of genes expressed in these cell cycle phases include tubulins, proteasome subunits, cyclins, centromere proteins, and histones, among others. The Zhang signature^25^, which was developed using cell cycle genes explicitly, is, as expected, correlated with all phases of the cell cycle. In contrast, the other signatures do not correlate with the cell cycle. When we compared the patient’s ColoTubIn score with the score of each cell cycle phase, we observed that the set of patients with high ColoTubIn clustered together (**Supplementary Figure S4 d-h**)

### 3.4 ColoTubIn association with tubulin inhibitors drugs

As shown in previous sections ColoTubIn is associated with the cell cycle and predicts survival of colon cancer patients. In order to test whether the ColoTubIn signature has not only prognostic but also predictive potential, we performed a drug-set enrichment analysis, mimicking gene set enrichment analysis (GSEA) (**Figure 1.6**). To do this we predicted the CPH probability, using the ColoTubIn CPH model in cell line expression data from the Cancer Cell Line Encyclopedia (CCLE) dataset^67^ and correlated it with the drug response from the PRISM drug repurposing dataset^68^. Based on the assumption that drugs sharing similar MoA produce comparable effects in cell lines, we performed a GSEA-like enrichment analysis linking drugs and MoA to the correlation between drug response and ColoTubIn CPH probability, while considering the directionality of the correlation. The most important hit was the MoA *tubulin polymerization inhibitor* (**Figure 5a**), which includes some of the drugs most strongly correlated with ColoTubIn CPH score (**Figure 5 b-e, Supplementary Figure S5 and S6**). Other significant MoAs for ColoTubIn are *bromodomain inhibitor,* and *EGFR inhibitor* drugs.

**Figure 5:**
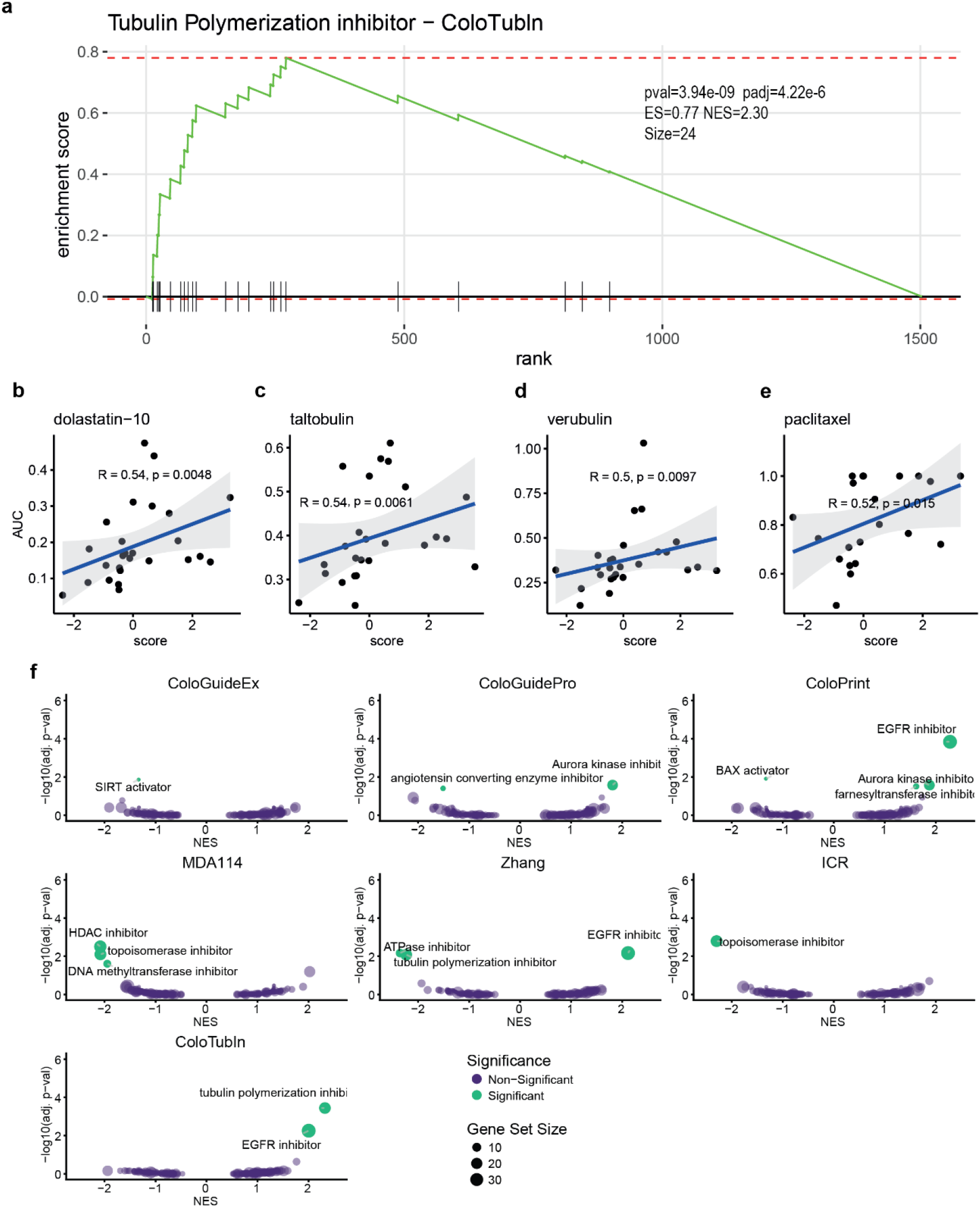
ColoTubIn is predictive of the response to tubulin inhibitor drugs. Panel a) shows the results of the enrichment between the ColoTubIn probability score and tubulin polymerization inhibitor (note that paclitaxel, which is a tubulin depolymerization inhibitor, is wrongly annotated as a polymerization inhibitor in PRISM) MoA as a GSEA plot. Most tubulin drugs cluster on the left of the plot, which represents the higher correlation with the ColoTubIn score. Panels b-e show the correlation between the ColoTubIn score and the top drugs from the tubulin inhibitor MAE. Panel f) shows all the enriched MAEs with normalized enrichment score (NES) on the x-axis and the -log10 p-value on the y-axis, for ColoTubIn and the other six signatures [Coloprint, MDA114, cologuideEx, coloGuidePro, Zhang and ICR]. Only the significant (p-value<0.05) MAEs are labeled. R represents the Pearson correlation.

Finally, we compared the main hits obtained from the “drug-set enrichment analysis” from ColoTubIn with the main hits obtained in the other signatures (**Figure 5f)**. The association of ColoTubIn with the tubulin *polymerization inhibitor* is statistically significant, whereas this was not observed in the other signatures, except for Zhang. These results indicate that ColoTubIn is specifically associated with tubulin inhibitor drugs. As expected, the Zhang signature, developed from genes implicated in cell cycle regulation, also displays enrichment for tubulin inhibitor drugs, however with a comparatively weaker association^25^. Consequently, this finding suggests that ColoTubIn has the potential to identify patients likely to benefit from this class of therapeutic agents.

### 3.5 Functional validation of sensitivity to Paclitaxel in preclinical models of colon cancer

In order to validate the ability of ColoTubIn to identify patients who would benefit from tubulin inhibitor treatment, we performed RNA-Sequencing on 80 PDX models obtained from colon cancer patients from the collection of Hector Palmer’s Group^64^. We then applied the ColoTubIn CPH model and predicted the five most resistant and the five most sensitive PDX models to tubulin inhibitors and consequently generated 10 corresponding PDXO models. Six PDXOs were selected based on their ability to efficiently grow as PDXO cultures: three PDXO models with a high ColoTubIn expression (CTAX23, CTAX26, and T138) and three with a low ColoTubIn expression (T224, T245, and T251) were chosen. Here, we tested *in vitro* sensitivity of the PDXO models to paclitaxel, a tubulin inhibitor that is widely used in other tumour types such as lung, breast, or ovarian cancer. We confirmed that all PDXO models with a high ColoTubIn probability were resistant to paclitaxel (**Figure 6a**). In contrast, the three PDXO models with a low ColoTubIn probability presented different therapeutic responses to paclitaxel. T224 and T251 models showed strong and moderate paclitaxel sensitivity, respectively. On the contrary, the T245 model presented resistance to paclitaxel (**Figure 6b**). The results were further evaluated using the maximum effect (Emax) and AUC, which have been proposed as suitable metrics for assessing paclitaxel response due to their strong association with cell viability. As expected, T224 showed the greatest sensitivity to paclitaxel according to both AUC and Emax, followed by T251, while the relative resistance of T245 was also confirmed (**Figure 6c**). CTAX23 displayed an unexpectedly low Emax; however, this estimate should be interpreted cautiously because the tested concentration range did not reach a clear response plateau, thereby limiting the reliable estimation of Emax ^69^.

**Figure 6:**
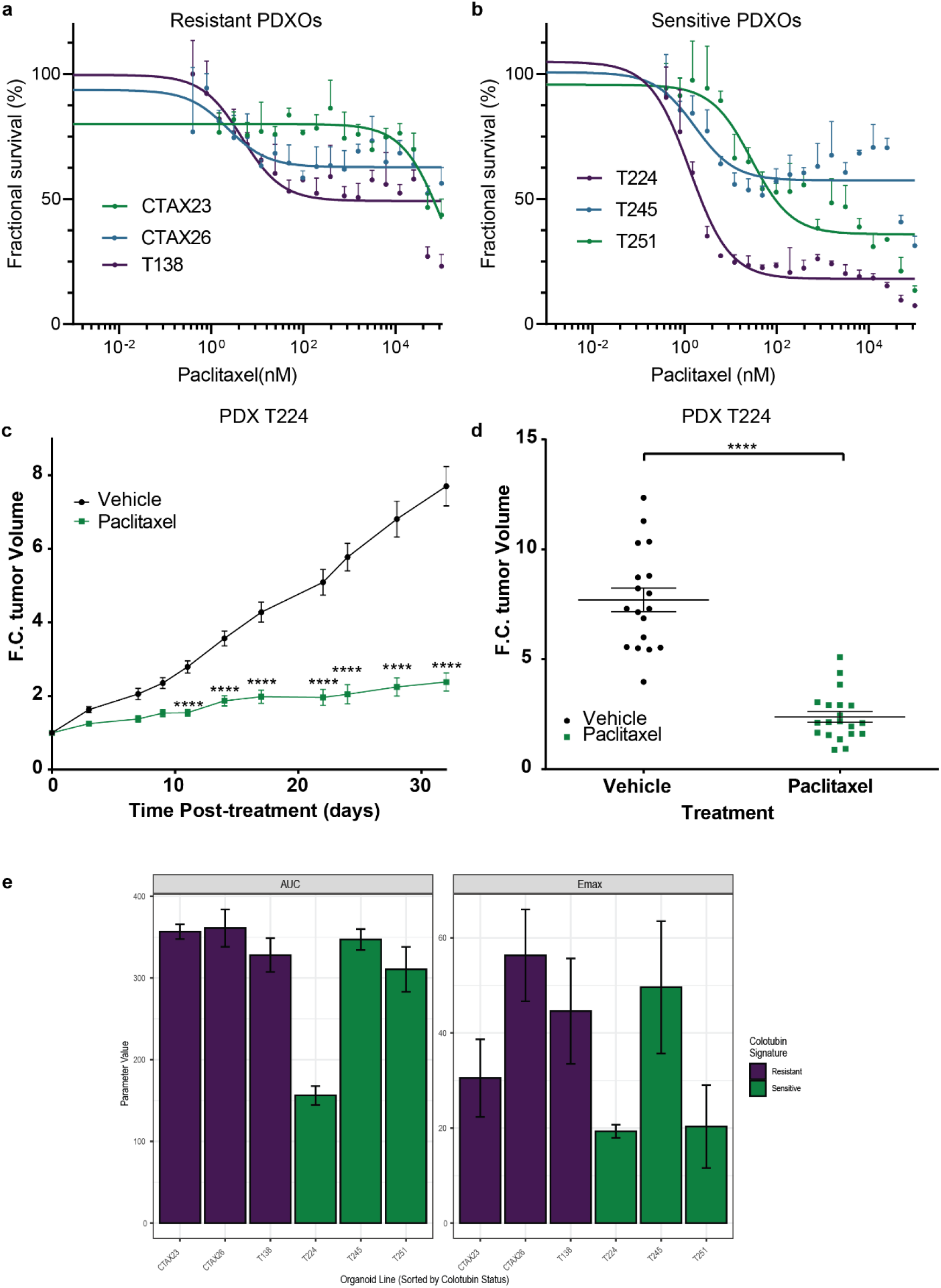
ColoTubIn treatment prediction to Paclitaxel sensitivity in vitro and in vivo validation. Panel a) shows dose–response curves of PDXOs CTAX23, CTAX26 and T138, identified as resistant by ColoTubIn to Paclitaxel. Panel b) illustrates dose-response curves of PDXOs T224, T245 and T251, identified as sensitive by ColoTubIn to Paclitaxel. Mean ± SD of triplicates is shown Panel c) show the drug response parameter AUC and Emax of resistant and sensitive PDXOs. Panel d) represent the relative tumour volume growth of T224 PDX (sensitive according to ColoTubIn) on treatment with vehicle (black) or Paclitaxel (green). Panel e) show a boxplot comparing relative tumour volume at day 32 of T224 PDX on treatment with vehicle (black) or Paclitaxel (green). Mean ± SEM is shown. Significant differences were assessed using one-way ANOVA and Dunnett’s multiple comparisons tests compared to vehicle group (****p value < 0.0001).

In order to explain the contrary result of T245 model (resistant when we predict sensitive), we performed a transcriptomic enrichment analysis in signatures of multi-drug resistance (ABC and SLC transporters)^70,71^, revealing that T245 preclinical model is highly enriched in the ABC transporters signature (**Figure S7a**), with one of the highest ABC transporter activities in the whole PDX cohort (8 out of 80 PDXs, **Figure S7b**). This result suggests that T245 is resistant not only to tubulin inhibitor drugs but also to many other drugs.

To validate our results, we used one of our sensitive preclinical colon cancer model (T224) to perform an *in vivo* experiment. First, we confirmed that the treatment with paclitaxel did not result in any significant weight reduction, or signs of toxicity compared with vehicle-treated NOD-SCID mice (data not shown). Mice bearing subcutaneous T224 tumours were treated intraperitoneally with vehicle (N=10) or paclitaxel (N=10). The results revealed a significant reduction in tumour growth in the paclitaxel-treated mice compared with the vehicle-treated mice (**Figure 6c-d**).

## 4. Discussion

In this work, we present a novel 37-gene expression-based signature for colon cancer with prognostic and predictive capabilities. Beyond demonstrating robust performance compared to 14 previously reported signatures in predicting overall survival, our model uniquely identifies a subgroup of colon cancer patients with shared molecular features who may benefit from tubulin inhibitors.

Validation *in vitro* and *in vivo* patient-derived preclinical models demonstrated the potential of the ColoTubIn signature in repurposing paclitaxel, a tubulin inhibitor. Additionally, interpretation of signature scores suggests that patients likely to benefit from these drugs tend to also have a poorer prognosis **(Figure 2c)**. These findings hold promise for further exploration in clinical trials.

A substantial innovation of this signature, driven by the methods used for its development, is its distinct, orthogonal biological information, which shows minimal correlation with other signatures or other clinical or molecular variables (**Figure 3c**). We examined our signature for enrichment in clinical variables and in the mutational status of relevant colon cancer genes, such as *APC*, *KRAS*, *TP53*, *BRAF*, and *SMAD4*. Importantly, the genes comprising the signature may not be direct drivers of the response mechanism or prognosis, but rather proxies for those factors. Consequently, the presented result emphasises that, unlike the compared signatures which largely recapitulate established molecular features, our signature shows more independence from known biomarkers. This underscores ColoTubIn’s promise as a new biomarker, largely distinct from existing ones.

Our findings indicate that the genes within ColoTubIn are strongly correlated with specific phases of the cell cycle, particularly with the mitotic phase (with inverse correlations of -0.6 and -0.66 with prophase and telophase-cytokinesis, respectively, according to Pearson correlation). This correlation elucidates the ability of ColoTubIn to predict patients who are likely to respond to tubulin inhibitor drugs, which disrupt microtubule dynamics during cell division. Our experimental findings highlight the robustness of ColoTubIn in identifying responsive patients.

Other microtubule targeting agents have been proposed for colon cancer treatment. For example, in Mertens et al^72^ several microtubule target agents, including vinca-alkaloids and taxanes, show significant responses in *KRAS*-mutated PDOs. Furthermore, plocabulin^65,73,74^ demonstrated activity in CRC patient-derived organoids (PDOs), and a phase II clinical trial has been completed (NCT03427268), which demonstrated disease stabilisation for over three months in 24,1% of participants^75^. *BRAF*(V600E) mutated or BRAF-like colon cancer has shown a vulnerability to vinorelbine^76^ and identified a synthetic lethal interaction between *BRAF*(V600E) and *RANBP2*, a member of the RAS family associated with mitosis and microtubule dynamics. Following these results, the MoTriColor clinical trial (EORTC1616, NCT03482362) was opened to further study vinorelbine activity in CRC, unfortunately was closed due to lack of enrolment. Generally, microtubule-targeting agents are not used to treat colon cancer because a clinical benefit could not sufficiently be demonstrated in clinical trials. For example, three trials on vinorelbine in colon cancer patients^77–79^ were negative. Nevertheless, the trial^79^ included 15 metastatic CRC patients of whom one patient (*BRAF* mutated) was in remission. A phase II trial of eribulin demonstrated limited anti-tumour activity in *BRAF*(V600E) mutated CRC patients, however some reduction in tumour size was observed^80^. Other studies using docetaxel showed also limited or no evidence of clinical benefit^81–83^. Therefore, it is crucial to restrict the use of tubulin-targeting agents to patients who are likely to benefit, as determined by biomarkers such as the one proposed in this study.

While our study yields meaningful insights, it is important to acknowledge its limitations. Although the efficacy of paclitaxel was tested in vitro and vivo, it needs to be highlighted that our cell line analysis presented similar or higher correlation with dolastatin, taltobulin, or verubulin. However, due to the availability of paclitaxel we focused our experimental validation on this tubulin inhibitor. Nevertheless, the use of predictive biomarkers for tubulin inhibitors facilitates the exploration of alternative drug treatment options, including dolastatin, taltobulin, or verubulin in potential future studies. Moreover, we acknowledge that ColoTubIn will only benefit a subgroup of patients. Yet, considering the promising outcomes achieved with current targeted therapies in BRAF-mutated colorectal cancer^7^, represented by a small subgroup of CRC patients, these findings underscore the value of developing therapies for defined patient subgroups. Furthermore, given the high prevalence of colon cancer, even a defined subset of patients may constitute a considerable population. Apart from this, it should be highlighted that in our case mapping probes to genes for signatures, which used different platforms, can introduce bias^84^. We also recognise that comparing stage-specific signatures (ColoPrint, ColoGuideEx and ColoGuidePro) with our TCGA-COAD cohort, which encompasses patients from all disease stages, presents certain challenges. Nonetheless, as our signature was derived from dataset including all clinical stages, it performed robustly across stage-specific comparisons. Next, we acknowledge that the in vitro validation of used a small, spectrum-enriched set of six PDXOs selected from extremes, which can limit generalisability. In a future study validation across a larger consecutive PDXO cohort is recommended.

Treating second-line and refractory colon cancer is an urgent unmet medical need, with no approved biomarker-based therapies available in the non-metastatic setting. The developed signature, when implemented in a test for paraffin-embedded samples, could offer clinicians a powerful new option for providing targeted treatment to these patients, which showed particular poor outcome in our analysis. A key advantage of paclitaxel is its extensively validated safety, tolerability, and side-effect profile, which supports rapid and cost-effective translation into clinical use.

In conclusion, our study revealed the prognostic and predictive potential of the ColoTubIn signature and highlights its unique potential of identifying colon cancer patients who respond to tubulin-targeting agent drugs. By demonstrating a strong correlation between ColoTubIn genes and the cell cycle, particularly the mitotic phase, we elucidated the biological basis for its predictive capacity, which we validated *in vitro* and *in vivo*. Our findings highlight the importance of considering ColoTubIn as a valuable tool in personalised medicine approaches for colon cancer treatment. ColoTubIn could transform the current management landscape in the second-line refractory disease and subsequently give new hope for a subset of patients diagnosed with a highly fatal malignancy.

## Data availability

The gene expression and related clinical data used in this study is publicly available via de GDC Data portal (https://portal.gdc.cancer.gov/) for TCGA datasets or via the GEO (https://www.ncbi.nlm.nih.gov/geo/) for GSE17536, GSE17537, GSE29621 and GSE39582 and via ArrayExpress under accession number for E-MTAB-12862 (Sarkans, U. et al. The BioStudies database—one stop shop for all data supporting a life sciences study. Nucleic Acids Res. 46, D1266–D1270 (2018). The SPINAL cohort was retrieved from SCORT database upon successful data request approval (https://www.s-cort.org).

## Code Availability

The code used for the analysis in this study is publicly available from the following repository: https://github.com/CancerCompBioLab/ColoTubIn. All the input files needed to replicate our findings, and the results obtained during the study are also available from the GitHub repository.

## Authors’ contributions

JLB, CFL, and JAS conceived the study design. JLB, LL, AMA, and IC analysed the data. JLB, LL, IC, JR, AMA, JMQ, EE, HGP, CFL, and JAS interpreted the data and results. JLB, LL, CFL, and JAS drafted the paper and figures. All authors critically reviewed the paper and the results and approved the final version.

## Acknowledgements

We first and foremost thank the participants, and their families included in the used studies. We are grateful to the investigators and data management teams who recruited the participants and to the pathologists who collected the samples. We thank the members of the Cancer Computational Biology Group for their helpful discussions. This work was supported by CMS2022-135428, RYC2019-026576-I, PID2020-115097RA-I00, AECC lab (LABAE259065SEOA) and ISCIII grant FORT23/00034 (to J.A. Seoane), Interreg Sudoe grant S1/1.1/P0033 co-financed by the European Regional Development Fund (ERDF) (to C. Fernandez-Lozano), and Postdoctoral fellowship from Galician Government ED481B_072 (to J.Liñares-Blanco).

## Supplementary Materials

### Supplementary Tables

**Supplementary Table S1:** P-values (transformed-ln(pvalue)) from statistical tests performed by comparing enrichment scores for each signature with clinical variables. The Wilcoxon test was performed on variables with two groups, whereas the Kruskal-Wallis test was used for those with three or more groups.

|  | **MDA114** | **coloGuideEx** | **coloPrint** | **ICR** | **Zhang** | **coloGuidePro** | **ColoTubIn** |
|---|---|---|---|---|---|---|---|
| **Consensus clusters** | 47.74 | 30.76 | 20.65 | 15.21 | 26.6 | 9.25 | 2.21 |
| **MSI** | 25.45 | 28.36 | 10.23 | 11.59 | 1.58 | 0.05 | 2.3 |
| **CIN vs GS** | 15.55 | 21.06 | 10.14 | 10.67 | 0.41 | 4.04 | 2.58 |
| **BRAF** | 15.64 | 13.36 | 8.92 | 7.5 | 0.19 | 0.98 | 1.89 |
| **APC** | 9.97 | 5.09 | 7.48 | 5.38 | 1.70 | 0.38 | 2.24 |
| **TP53** | 4.73 | 2.41 | 2.15 | 0.91 | 0.48 | 4.18 | 1.89 |
| **SMAD4** | 1.57 | 1.37 | 0.24 | 0.39 | 1.26 | 0.95 | 1.51 |
| **KRAS** | 0.31 | 0.83 | 1.07 | 1.43 | 0.05 | 1.73 | 1.12 |

**Supplementary Table S2:** Clinical characteristics across discovery and validation cohorts.

| **Variable** | **Overall** N = 1,695<sup>*1*</sup> | **TCGA** N = 325<sup>*1*</sup> | **EMTAB12862** N = 782<sup>*1*</sup> | **GSE17536** N = 177<sup>*1*</sup> | **GSE17537** N = 55<sup>*1*</sup> | **GSE29621** N = 65<sup>*1*</sup> | **GSE39582** N = 185<sup>*1*</sup> | **SPINAL** N = 106<sup>*1*</sup> |
|---|---|---|---|---|---|---|---|---|
| **Age at Diagnosis** | 70 (61, 78) | 68 (58, 78) | 74 (65, 81) | 66 (57, 75) | 62 (54, 72) | NA (NA, NA) | 64 (54, 71) | 74 (65, 83) |
| Missing | 65 | 0 | 0 | 0 | 0 | 65 | 0 | 0 |
| **Survival Time (Days - 5yr Capped)** | 1,598 (697, 1,825) | 596 (362, 1,074) | 1,825 (1,376, 1,825) | 1,268 (683, 1,825) | 1,506 (925, 1,801) | 1,434 (671, 1,799) | 1,680 (870, 1,825) | 1,412 (751, 1,825) |
| **Vital Status (at 5 yrs)** |  |  |  |  |  |  |  |  |
| Alive | 1,188 (70%) | 253 (78%) | 537 (69%) | 110 (62%) | 35 (64%) | 42 (65%) | 141 (76%) | 70 (66%) |
| Dead | 507 (30%) | 72 (22%) | 245 (31%) | 67 (38%) | 20 (36%) | 23 (35%) | 44 (24%) | 36 (34%) |
| **Gender** |  |  |  |  |  |  |  |  |
| Female | 823 (49%) | 160 (49%) | 405 (52%) | 81 (46%) | 29 (53%) | 25 (38%) | 80 (43%) | 43 (41%) |
| Male | 872 (51%) | 165 (51%) | 377 (48%) | 96 (54%) | 26 (47%) | 40 (62%) | 105 (57%) | 63 (59%) |
| **Tumor Stage** |  |  |  |  |  |  |  |  |
| Stage 1 | 199 (12%) | 54 (17%) | 90 (12%) | 24 (14%) | 4 (7.3%) | 7 (11%) | 0 (0%) | 20 (19%) |
| Stage 2 | 616 (36%) | 125 (38%) | 326 (42%) | 57 (32%) | 15 (27%) | 22 (34%) | 47 (25%) | 24 (23%) |
| Stage 3 | 614 (36%) | 99 (30%) | 287 (37%) | 57 (32%) | 19 (35%) | 18 (28%) | 109 (59%) | 25 (24%) |
| Stage 4 | 266 (16%) | 47 (14%) | 79 (10%) | 39 (22%) | 17 (31%) | 18 (28%) | 29 (16%) | 37 (35%) |
| **MSI Status** |  |  |  |  |  |  |  |  |
| MSI | 442 (26%) | 57 (18%) | 217 (28%) | 57 (32%) | 27 (49%) | 13 (20%) | 51 (28%) | 20 (19%) |
| MSS | 1,253 (74%) | 268 (82%) | 565 (72%) | 120 (68%) | 28 (51%) | 52 (80%) | 134 (72%) | 86 (81%) |

<sup>*1*</sup>Median (Q1, Q3); n (%)

**Supplementary Table S3:**
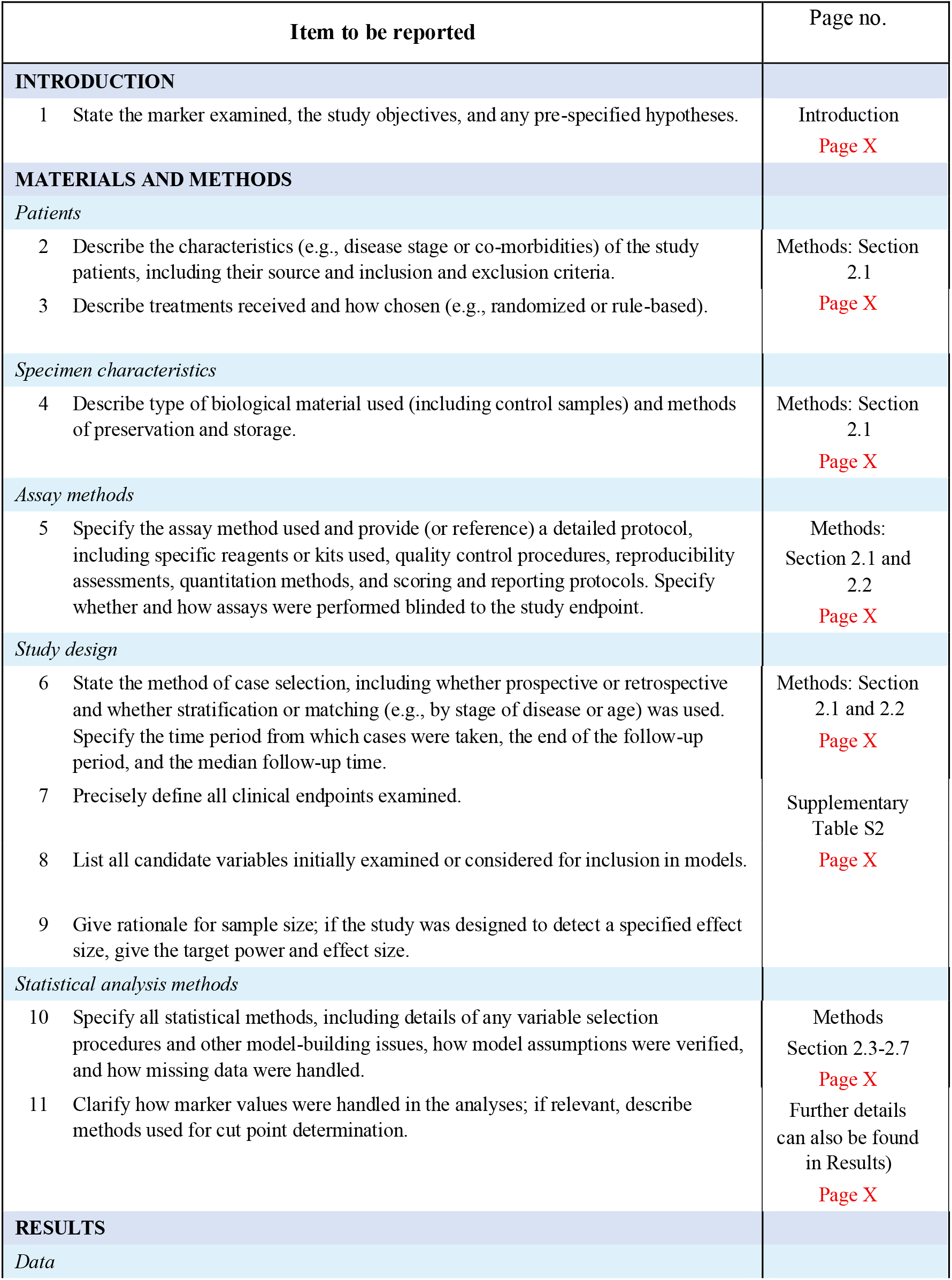
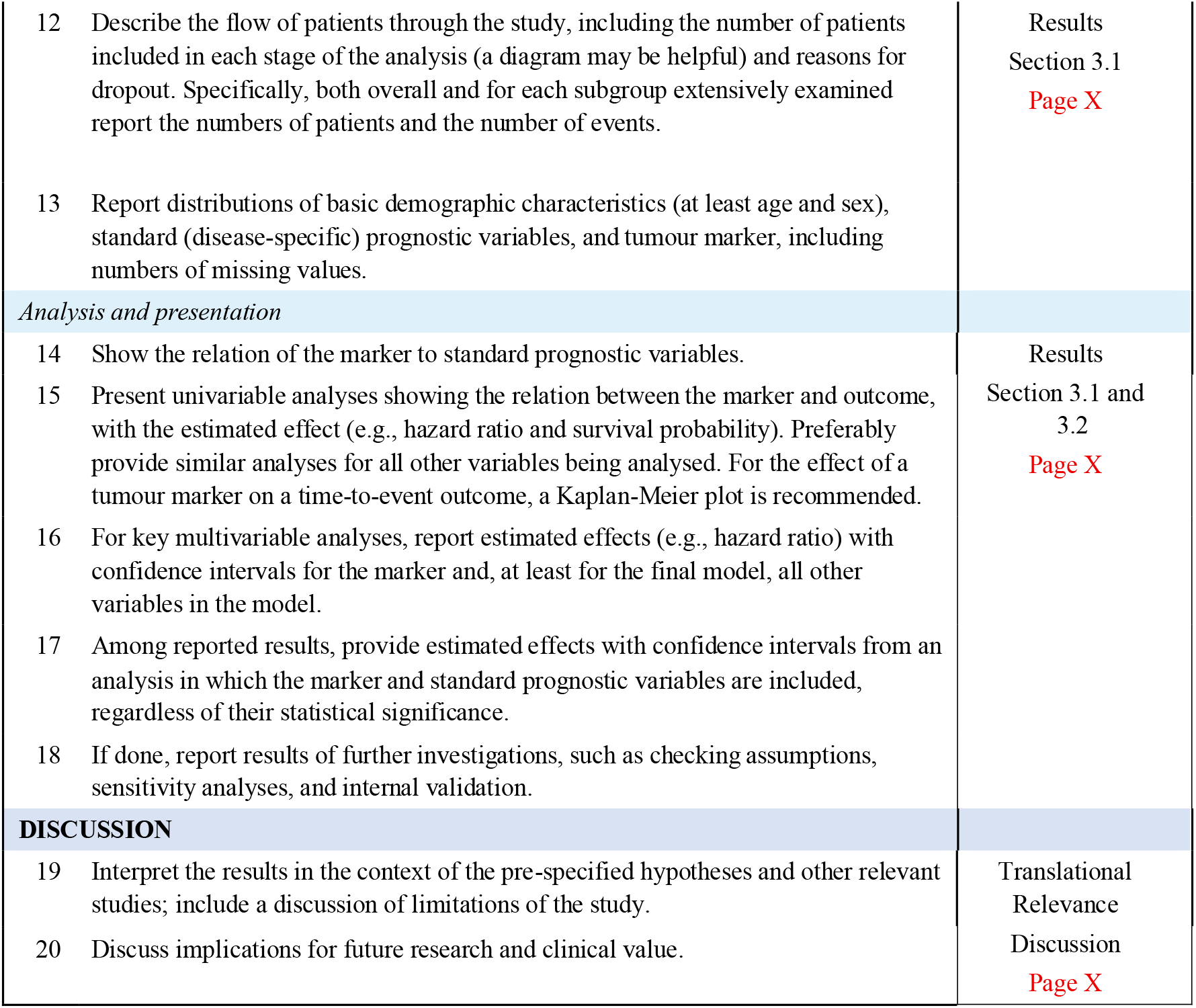
REMARK criteria checklist.

| Item to be reported |  | Page no. |
|---|---|---|
| **INTRODUCTION** |  |  |
| 1 | State the marker examined, the study objectives, and any pre-specified hypotheses. | Introduction<br>Page X |
| **MATERIALS AND METHODS** |  |  |
| *Patients* |  |  |
| 2 | Describe the characteristics (e.g., disease stage or co-morbidities) of the study patients, including their source and inclusion and exclusion criteria. | Methods: Section 2.1<br>Page X |
| 3 | Describe treatments received and how chosen (e.g., randomized or rule-based). |  |
| *Specimen characteristics* |  |  |
| 4 | Describe type of biological material used (including control samples) and methods of preservation and storage. | Methods: Section 2.1<br>Page X |
| *Assay methods* |  |  |
| 5 | Specify the assay method used and provide (or reference) a detailed protocol, including specific reagents or kits used, quality control procedures, reproducibility assessments, quantitation methods, and scoring and reporting protocols. Specify whether and how assays were performed blinded to the study endpoint. | Methods:<br>Section 2.1 and 2.2<br>Page X |
| *Study design* |  |  |
| 6 | State the method of case selection, including whether prospective or retrospective and whether stratification or matching (e.g., by stage of disease or age) was used. Specify the time period from which cases were taken, the end of the follow-up period, and the median follow-up time. | Methods: Section 2.1 and 2.2<br>Page X |
| 7 | Precisely define all clinical endpoints examined. | Supplementary Table S2<br>Page X |
| 8 | List all candidate variables initially examined or considered for inclusion in models. |  |
| 9 | Give rationale for sample size; if the study was designed to detect a specified effect size, give the target power and effect size. |  |
| *Statistical analysis methods* |  |  |
| 10 | Specify all statistical methods, including details of any variable selection procedures and other model-building issues, how model assumptions were verified, and how missing data were handled. | Methods<br>Section 2.3-2.7<br>Page X |
| 11 | Clarify how marker values were handled in the analyses; if relevant, describe methods used for cut point determination. | Further details can also be found in Results)<br>Page X |
| **RESULTS** |  |  |
| *Data* |  |  |
| 12 | Describe the flow of patients through the study, including the number of patients included in each stage of the analysis (a diagram may be helpful) and reasons for dropout. Specifically, both overall and for each subgroup extensively examined report the numbers of patients and the number of events. | Results<br>Section 3.1<br>Page X |
| 13 | Report distributions of basic demographic characteristics (at least age and sex), standard (disease-specific) prognostic variables, and tumour marker, including numbers of missing values. |  |
| *Analysis and presentation* |  |  |
| 14 | Show the relation of the marker to standard prognostic variables. | Results<br>Section 3.1 and 3.2<br>Page X |
| 15 | Present univariable analyses showing the relation between the marker and outcome, with the estimated effect (e.g., hazard ratio and survival probability). Preferably provide similar analyses for all other variables being analysed. For the effect of a tumour marker on a time-to-event outcome, a Kaplan-Meier plot is recommended. |  |
| 16 | For key multivariable analyses, report estimated effects (e.g., hazard ratio) with confidence intervals for the marker and, at least for the final model, all other variables in the model. |  |
| 17 | Among reported results, provide estimated effects with confidence intervals from an analysis in which the marker and standard prognostic variables are included, regardless of their statistical significance. |  |
| 18 | If done, report results of further investigations, such as checking assumptions, sensitivity analyses, and internal validation. |  |
| **DISCUSSION** |  |  |
| 19 | Interpret the results in the context of the pre-specified hypotheses and other relevant studies; include a discussion of limitations of the study. | Translational Relevance<br>Discussion<br>Page X |
| 20 | Discuss implications for future research and clinical value. |  |

### Supplementary Figures

**Supplementary Figure S1:**
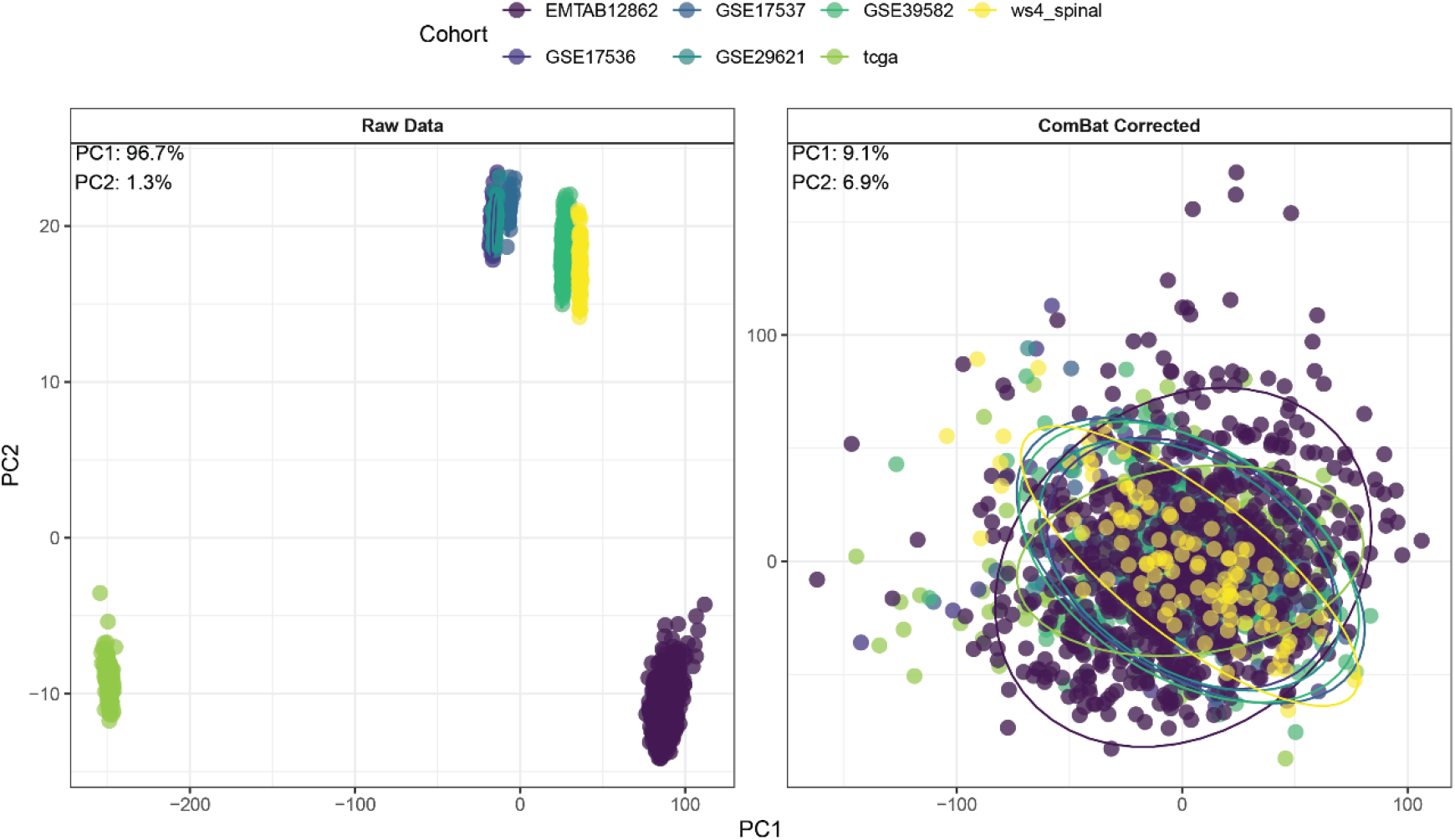
The left panel represents a scatter plot with the first two PCA before ComBat. The right panel shows the distribution after the elimination of the batch effect. Before the validation of the models, the batch effect between the cohorts was eliminated. For this purpose, the ComBat function of the SVA package was run, with the TCGA cohort samples used as the reference batch. The figure on the right shows homogeneity in the distribution of samples between cohorts.

**Supplementary Figure 2:**
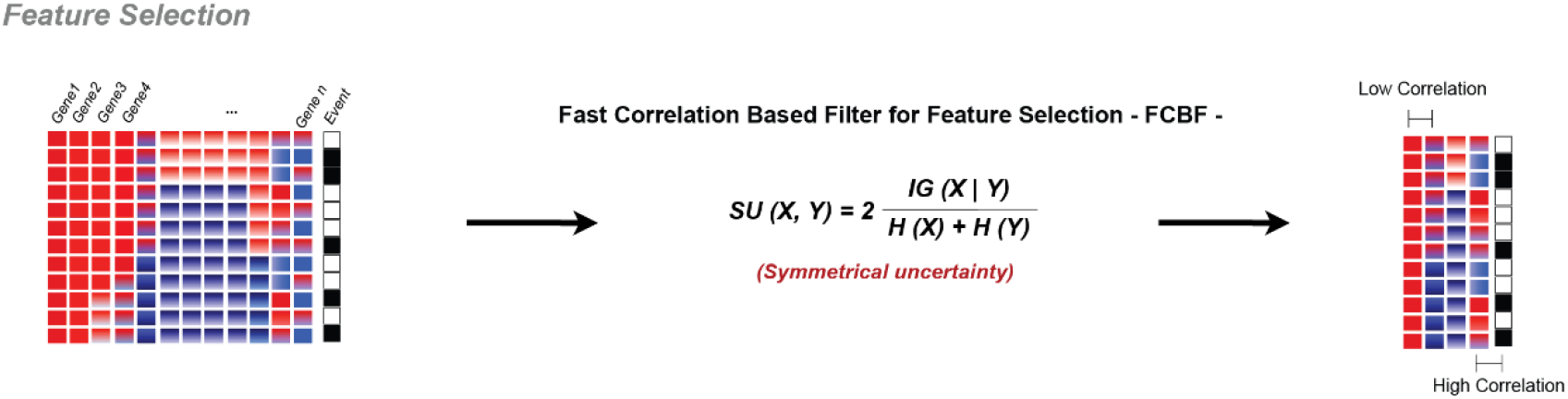
Schematic of how a Fast Correlation Based Filter (FCBF) was used to select features correlated with the outcome but not between them.

**Supplementary Figure S3:**
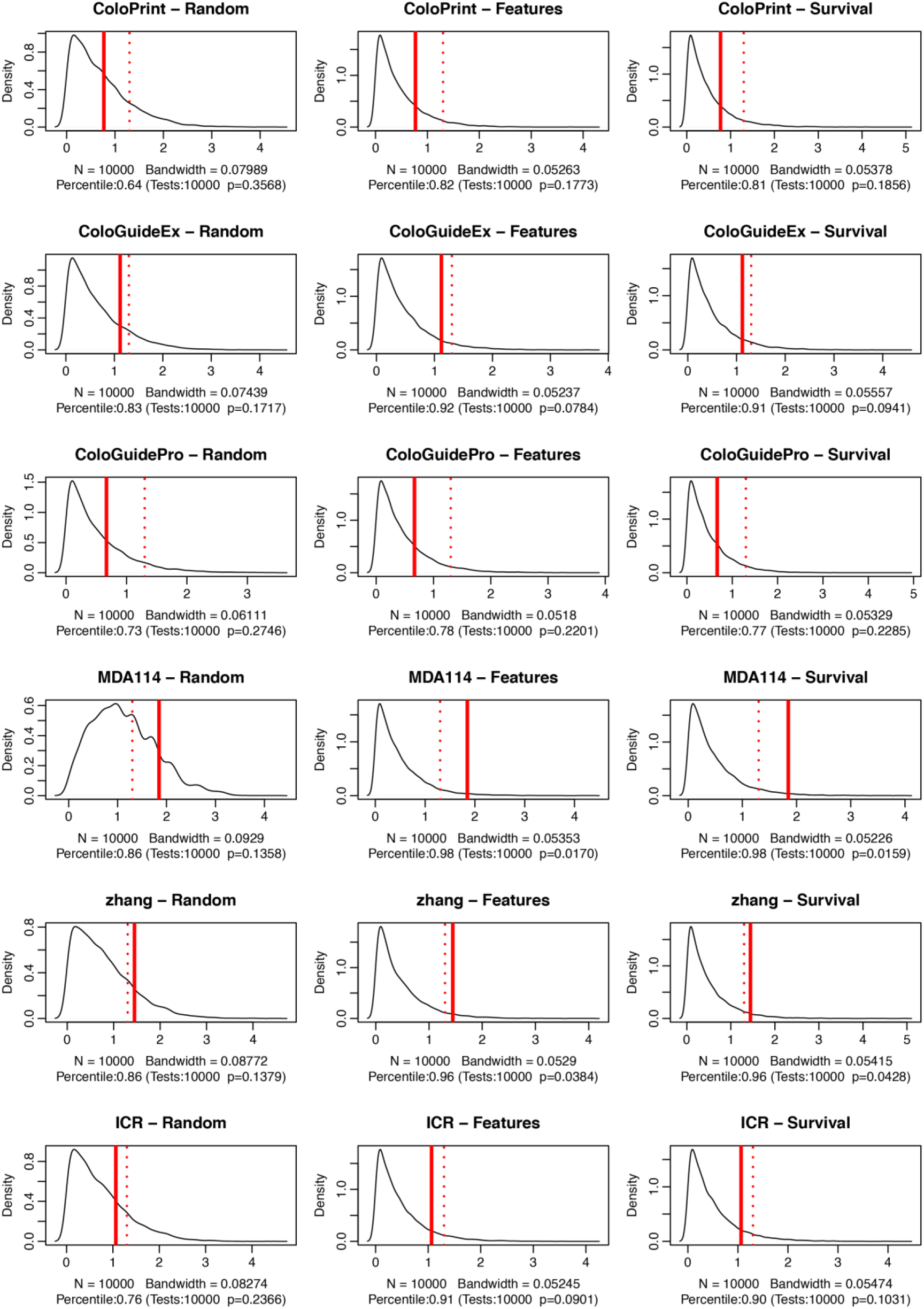
SigCheck validation of the coloPrint, coloGuideEx, coloGuidePro, MDA114, Zhang and ICR signatures.

**Supplementary Figure S4:**
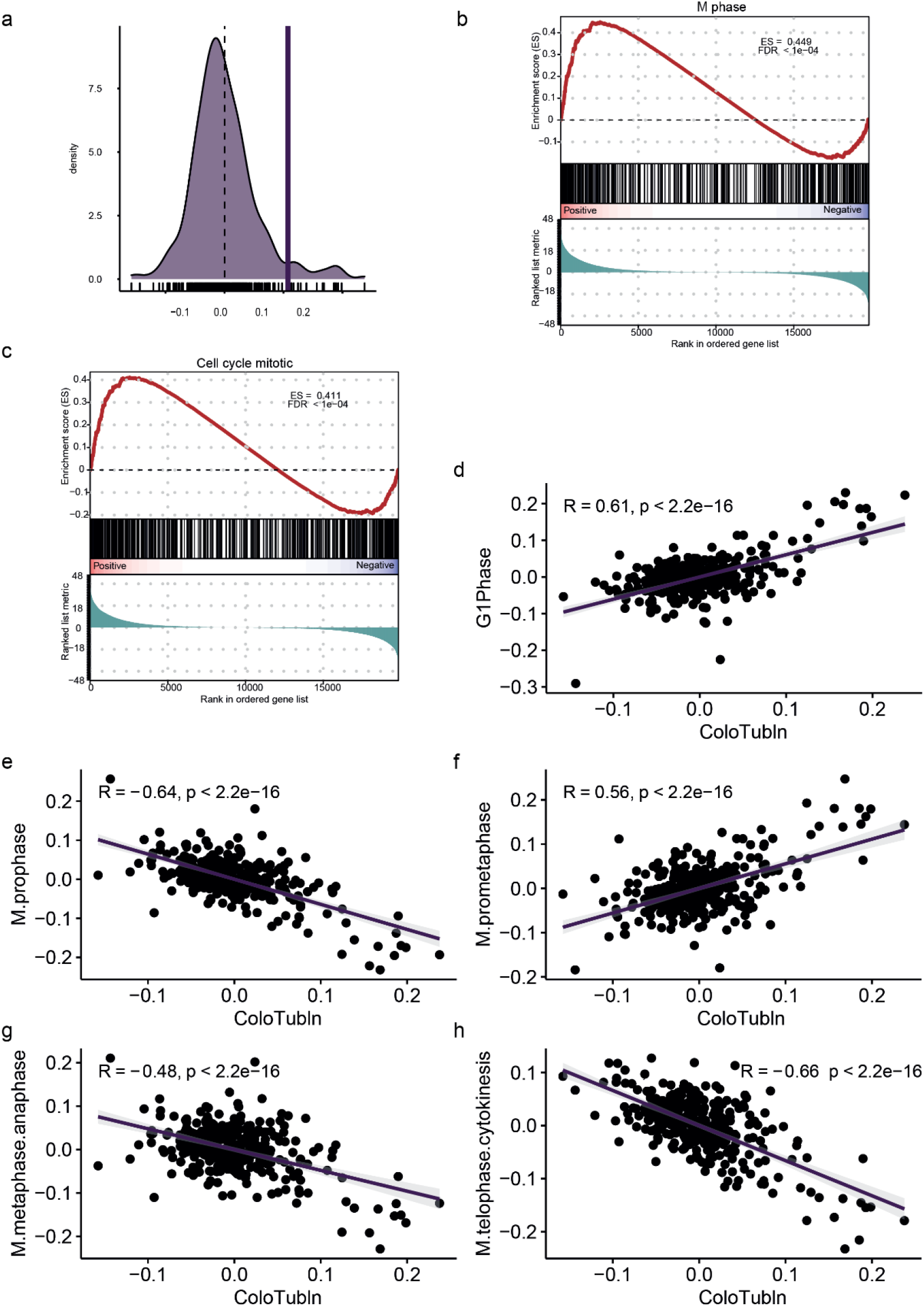
ColoTubIn pathway enrichment. a) Distribution of ColoTubIn score. The vertical line represents the separation between the low and high ColoTubIn activity clusters, defined by MClust. b) GSEA plot of M-phase gene set enrichment in high vs low ColoTubIn samples. c) GSEA plot of cell cycle mitotic gene set enrichment of high vs low ColoTubIn samples. Scatter plots showing the correlation of the ColoTubIn signature with cell cycle phases: d) G1-phase, e) M-prophase, f) M-prometaphase, g) M-metaphase-anaphase and h) M-telophase cytokinesis. R represents the Pearson correlation.

**Supplementary Figure S5:**
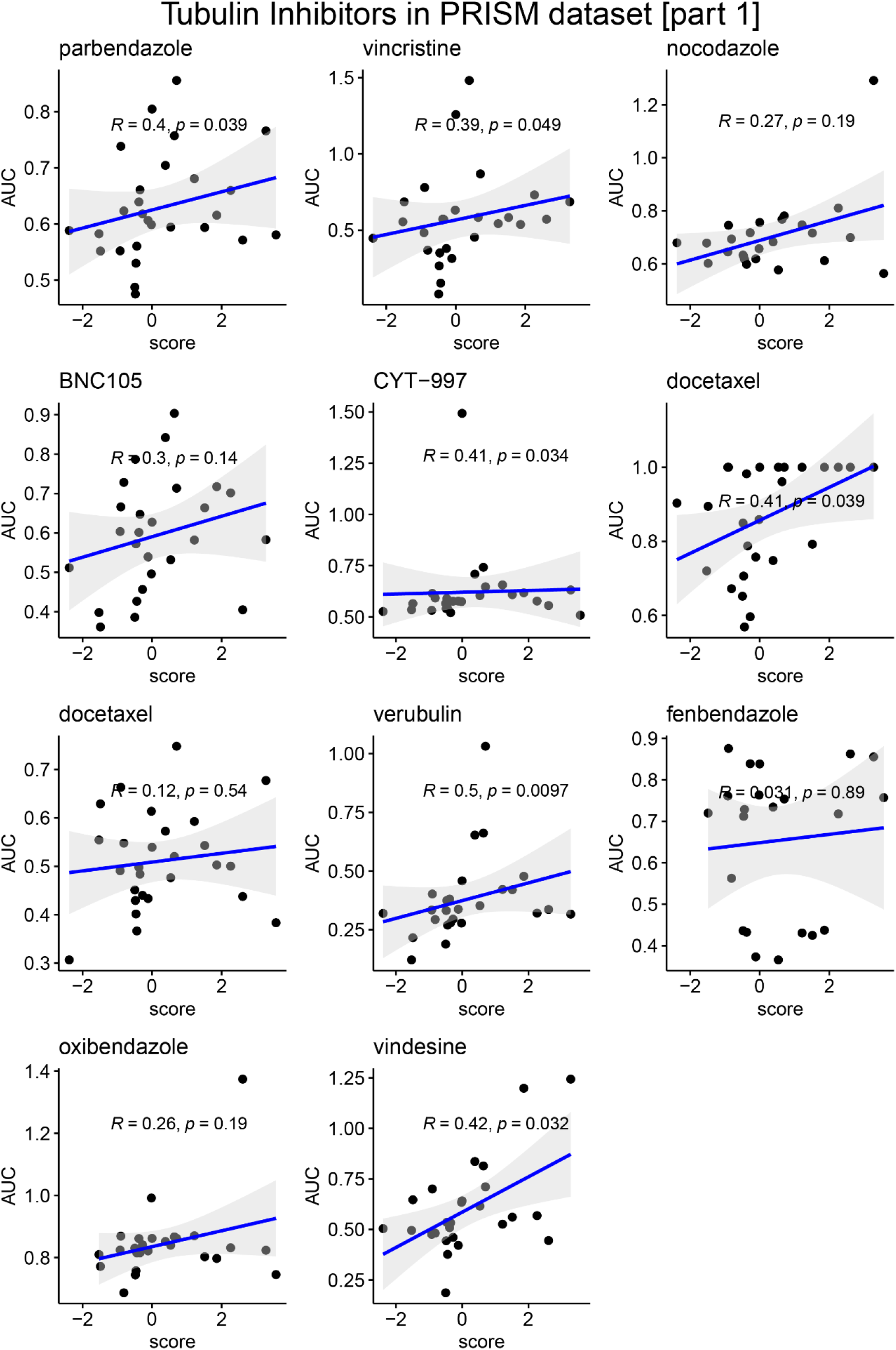
Scatter plots showing the ColoTubIn outcome prediction and tubulin inhibitor drug correlation (Part 1). R represents the Pearson correlation.

**Supplementary Figure S6:**
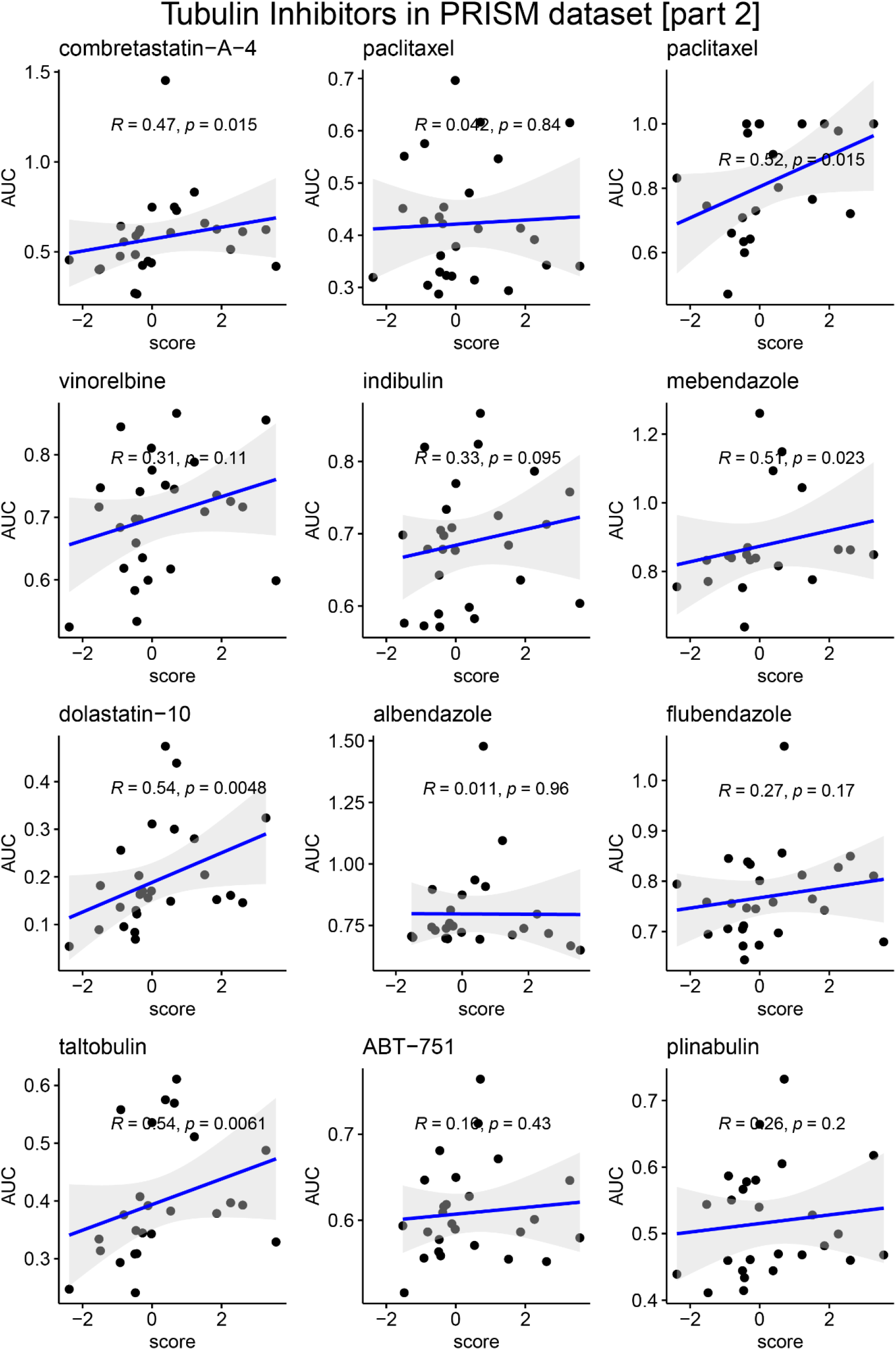
Scatter plots showing the ColoTubIn outcome prediction and tubulin inhibitor drug correlation (Part 2). R represents the Pearson correlation.

**Supplementary Figure S7:**
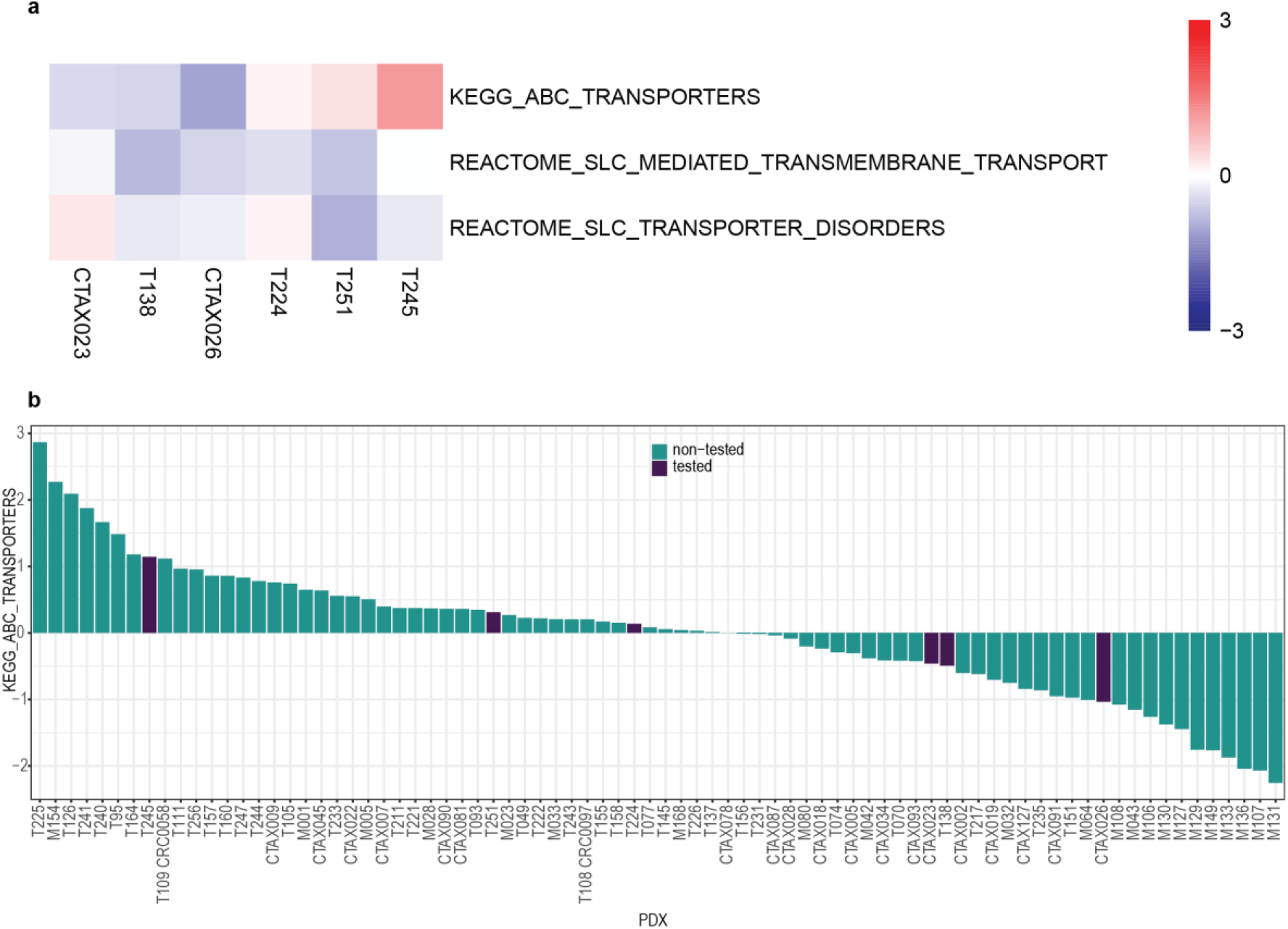
a) Enrichment of tested PDXOs in multi-drug resistance pathways from KEGG and REACTOME. T245 has the greatest enrichment in ABC transporters pathway activity b) Bar plot showing the enrichment of the ABC transporters signature in the PDX cohort (PDX sorted by signature enrichment). T245 has one of the highest activities among ABC transporters (8/80).

